# Modulation of peptidoglycan synthesis by recycled cell wall tetrapeptides

**DOI:** 10.1101/771642

**Authors:** Sara B. Hernández, Tobias Dörr, Matthew K. Waldor, Felipe Cava

**Affiliations:** The laboratory for Molecular Infection Medicine Sweden (MIMS), Department of Molecular Biology. Umeå University. Umeå. Sweden; Weill Institute for Cell and Molecular Biology, Department of Microbiology and Cornell Institute of Host-Microbe Interactions and Disease, Cornell University, Ithaca, NY 14853; Howard Hughes Medical Institute, Brigham and Women’s Hospital Division of Infectious Diseases and Harvard Medical School Department of Microbiology, Boston, Massachusetts, USA

**Keywords:** cell wall, *Vibrio cholerae*, peptidoglycan recycling, L,D-carboxypeptidase, L,D-transpeptidase, NCDAA, D-amino acids, Tn-seq

## Abstract

The bacterial cell wall is made of peptidoglycan (PG), a polymer that is essential for maintenance of cell shape and survival. During growth, bacteria remodel their PG, releasing fragments that are predominantly re-internalized by the cell, where they are recycled for synthesis of new PG. Although the PG recycling pathway is widely conserved, its components are not essential and its roles in cell wall homeostasis are not well-understood. Here, we identified LdcV, a *Vibrio cholerae* L,D-carboxypeptidase that cleaves the terminal D-Alanine from recycled murotetrapeptides. In the absence of *ldcV*, recycled tetrapeptides accumulated in the cytosol, leading to two toxic consequences for the cell wall. First, incorporation of tetrapeptide-containing PG precursors into the cell wall led to reduction in D,D-cross-linkage between stem peptides, diminishing PG integrity. Second, tetrapeptide accumulation led to a decrease in canonical UDP-pentapeptide precursors, reducing PG synthesis. Thus, LdcV and the recycling pathway promote optimal cell wall assembly and composition. Furthermore, Ldc substrate preference for murotetrapeptides containing canonical (D-Alanine) vs. non-canonical (D-Methionine) D-amino acids is conserved, suggesting that accumulation of tetrapeptide recycling intermediates may modulate PG homeostasis in environments enriched in non-canonical-muropeptides via substrate competition.

## INTRODUCTION

The bacterial cell shape-determining peptidoglycan (PG) cell wall, provides resistance to cell turgor pressure and protection from environmental threats^1, 2^. PG is a heteropolymer made up of glycan chains consisting of *N*-acetylglucosamine (GlcNAc) and *N*-acetylmuramic acid (MurNAc). The GlcNAc-MurNAc polymers are cross-linked via short peptides, forming a cell-size mesh known as the murein sacculus^1^. The composition of these peptides normally includes a terminal D-Alanine (D-Ala), however this amino acid is replaced in some species by non-canonical D-amino acids (NCDAAs)^3–5^ when cells enter stationary phase.

During growth, the PG sacculus expands through the coordinated action of degradative and synthetic enzymes^6, 7^. In *Escherichia coli*, 50% of the murein sacculus is thought to be cleaved at each generation^8^. Although some of the cleaved PG fragments are released into the extracellular medium^9–11^, most of them are transported back to the cytoplasm for their reutilization, a process referred to as the PG recycling pathway^12^. Cleavage of the sacculus by endopeptidases (EPs) and lytic transglycosylases (LTs) releases monomeric 1,6-anhydro-muropeptides^7^ that are specifically imported into the cytoplasm by the AmpG permease^13, 14^ to serve as a substrate for the NagZ β-*N*-acetylglucosaminidase^15, 16^ and the AmpD amidase^17, 18^ (**Fig. 1a**). The resulting products of these two enzymes are further cleaved by the L,D-carboxypeptidase LdcA into tripeptides^19^. The tripeptide is next transformed into UDP-MurNAc-tripeptide by the muropeptide ligase Mpl^20^, thereby connecting the PG recycling and *de novo* synthesis pathways (**Fig. 1a**).

**Fig. 1.**
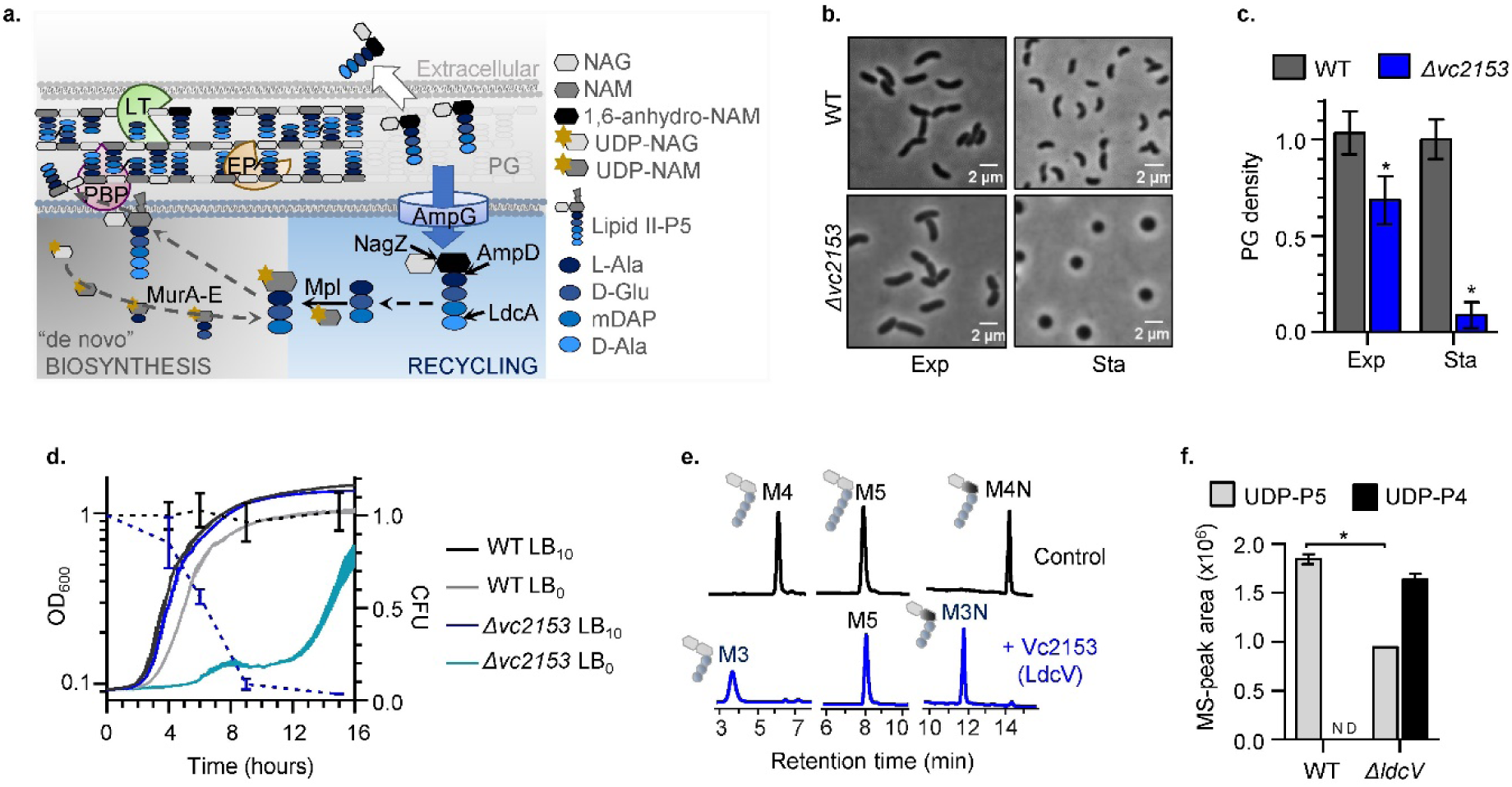
*vc2153* encodes an L,D-carboxypeptidase (LdcV) that contributes to peptidoglycan recycling in *V. cholerae*. **a.** Schematic representation of the PG recycling pathway in Gram negative bacteria (LT: lytic transglycosilase; EP: endopeptidase; PBP: penicillin binding protein). **b.** Phase contrast microscopy of exponential (Exp) or stationary (Sta) phase LB_10_ cells imaged on agarose pads. **c-d.** (**c**) PG density (normalised to the value of the WT), (**d**) growth kinetics of strains growing in LB containing 0 (LB_0_) or 10 (LB_10_) g/L of NaCl (left Y axis) and number of CFU (normalised to wt) counted in LB_10_ cultures at indicated times (dashed lines, right Y axis). **e.** Hydrolytic activity of purified Vc2153 protein tested on isolated monomeric muropeptides (M4: murotetrapeptide; M3: murotripeptide; M5: muropentapeptide; N: anhydro form of the muropeptide). **f.** Quantification by MS^e^ of UDP-muramyl-pentapeptide (UDP-P5) and UDP-muramyl-tetrapeptide (UDP-P4) detected in the cytosol of the indicated strains. Data represent mean ± s.d. of triplicates; ND: not detected; \**P* < 0.05, unpaired t-test.

Although PG recycling pathways are broadly conserved, this process of salvaging murein components is surprisingly not essential for bacterial growth^12^, at least under standard laboratory conditions. Only the absence of the cytoplasmic LdcA is lethal when *E. coli* enters into stationary growth phase; Höltje and colleagues suggested that the incorporation of atypical tetrapeptide PG precursors into the murein sacculus might result in a lethal cross-linkage defect because these muropeptides can only function as acceptors and not as donors in the cross-linking reaction^19^. However, incorporation of recycled tetrapeptide precursors into PG has not been demonstrated and the consequences of accumulation of these atypical precursors have not been explored.

Here we found that *V. cholerae vc2153* (designated LdcV), is a functional analogue of LdcA and encodes an L,D-carboxypeptidase. A *ΔlcdV* mutant strain exhibited a marked reduction in PG content, hypersensitivity to conditions of low-osmolarity and spherical morphology in stationary phase. Genetic analyses revealed that *ldcV* plays a critical role in the regulatory network controlling PG homeostasis. The tetrapeptides that accumulate in the *lcdV* mutant were found to be incorporated into the murein, which consequently becomes deficient in cross-links as predicted. Moreover, we found that accumulation of tetrapeptides coincides with a decrease in the levels of canonical UDP-P5 precursors, which leads to a reduction in PG synthesis. Finally, we demonstrate that recycling tetrapeptides accumulate in wild-type *V. cholerae* during stationary phase due to LdcV’s substrate preference for canonical (D-Ala) vs non-canonical (D-Met) containing murotetrapeptides, a property that is conserved in other Ldcs. Collectively our findings reveal that PG recycling and L,D-carboxypeptidases in particular exert a critical role in PG homeostasis, modulating PG synthesis and composition.

## RESULTS

### *V. cholerae vc2153* encodes an L,D-carboxypeptidase involved in peptidoglycan recycling

In a transposon screen to identify genes that interact with *mrcA*^21^, *V. cholerae’s* major PG synthetase (PBP1A), we found that *vc2153* was synthetically lethal with *mrcA*. Similar to a *mrcA* mutant^5, 22^, an unmarked in-frame deletion mutant of *vc2153* (*Δvc2153*) was spherical in stationary phase and had a marked reduction in PG content (**Fig. 1b-c**). However, the *Δvc2153* mutant exhibited a more severe reduction in viability that was particularly aggravated under low osmolarity conditions (salt free LB medium, LB0) (**Fig. 1d and Supplementary Fig. 1a**). Together these observations suggested that although VC2153 likely contributes to PG biogenesis, it does not appear to act in the PBP1A pathway.

VC2153 is annotated as a VanY D,D-carboxypeptidase-like protein in the NCBI database. However, using protein domain analysis^23^, we predicted that *vc2153* encodes a putative LdcB (L,D-carboxypeptidase). Bacterial carboxypeptidases are PG hydrolytic enzymes that remove the C-terminal amino acid from stem peptides of muropeptides^7^. While D,D-carboxypeptidases act on pentapeptides^24^, L,D-carboxypeptidases use tetrapeptides as substrates^25^. VC2153 was purified to test its carboxypeptidase activity using different monomeric muropeptides as potential substrates (**Fig. 1e**). Purified LdcA, the *E.coli* L,D-carboxypeptidase^19, 26^ was used as a positive control in these assays. VC2153 fully converted the disaccharide-tetrapeptide substrates (both M4 and its anhydro-derivative, M4N) to disaccharide-tripeptides (M3 and M3N), cleaving the peptide bond between *m*DAP and the terminal D-Ala. VC2153, like LdcA, did not act on pentapeptides (**Fig. 1e**) or cross-linked muropeptides (e.g. D44) (**Supplementary Fig. 1b**). Interestingly, in contrast to a previous report^26^, both LdcV and LdcA also acted on high molecular weight PG (**Supplementary Fig. 1c-e**), suggesting a potential role for L,D-carboxypeptidases in the modification of the sacculus. Together, these observations demonstrate that *V. cholerae* VC2153 (re-named LdcV) is an L,D-carboxypeptidase.

VCA0337 and VCA0439^19^, two proteins that differ in a single amino acid, and which have been implicated in microcin immunity^27, 28^, exhibit greater similarity to LdcA than LdcV (**Supplementary Fig. 1f**). However, purified VCA0337 had no PG-hydrolytic activity (**Supplementary Fig. 1g-h**). Furthermore, heterologous expression of *E. coli ldcA* under control of the PBAD promoter complemented the *ΔldcV* strain (**Supplementary Fig. 1i**), suggesting that LdcV is the functional homologue of *E. coli*’s L,D-carboxypeptidase LdcA.

LdcA plays a crucial role in the *E. coli* PG recycling pathway, converting soluble PG-tetrapeptides into tripeptides, which are then re-used for PG synthesis. The PG-tetrapeptides that accumulate in LdcA mutants are activated by Mpl to UDP-MurNAc-tetrapeptides (UDP-P4)^19, 29^. In contrast to UDP-P3 however, UDP-P4 is not a substrate of MurF, the ligase that produces the PG-pentapeptide UDP-P5 ^12^. Since pentapeptides, and not tetrapeptides, are donor substrates of D,D-transpeptidases^1^ (i.e., PBP TPases), it has been suggested that the lytic phenotype observed in the *ldcA* mutant results from disruption of PG-cross-linkage homeostasis^19^. To determine whether LdcV, like LdcA, is involved in PG recycling, we compared the pool of cytosolic UDP-activated murein precursors in the Δ*ldcV* mutant and the wild-type strain, using untargeted UPLC-MS. High levels of UDP-P4 were detected in the *ΔldcV* mutant, but were undetectable in the wild-type strain (**Fig. 1f and Supplementary Fig. 2**), supporting the idea that LdcV is involved in the *V. cholerae* PG recycling pathway.

### Phenotypic consequences of *ΔldcV* are associated with accumulation of UDP-MurNAc-tetrapeptide and cellular autolysis

Colonies of the *ldcV* mutant were visibly distinct from wild-type *V. cholerae* colonies on LB10 agar plates (**Fig. 2a**). However, after 3 days of growth on plates, wild-type appearing colonies emerged from the mutant colonies (**Fig. 2a and Supplementary Fig. 3a**). The reversion toward wild-type morphology was recapitulated upon colony re-isolation, suggesting that stable suppressors of Δ*ldcV* had developed. Additional analyses of 13 *ΔldcV* suppressor colonies (Sup1-13) revealed that they all at least partially restored wild-type phenotypes in cell morphology, PG density and growth in low osmolarity medium (**Fig. 2b and Supplementary Fig. 3b-c**). Remarkably, whole genome sequence analysis identified single-nucleotide polymorphisms (SNPs), primarily creating truncations, in genes that participate in the PG recycling pathway in all 13 strains (**Supplementary Fig. 3d-e**). These observations strongly suggest that PG recycling becomes toxic in the absence of LdcV.

**Fig. 2.**
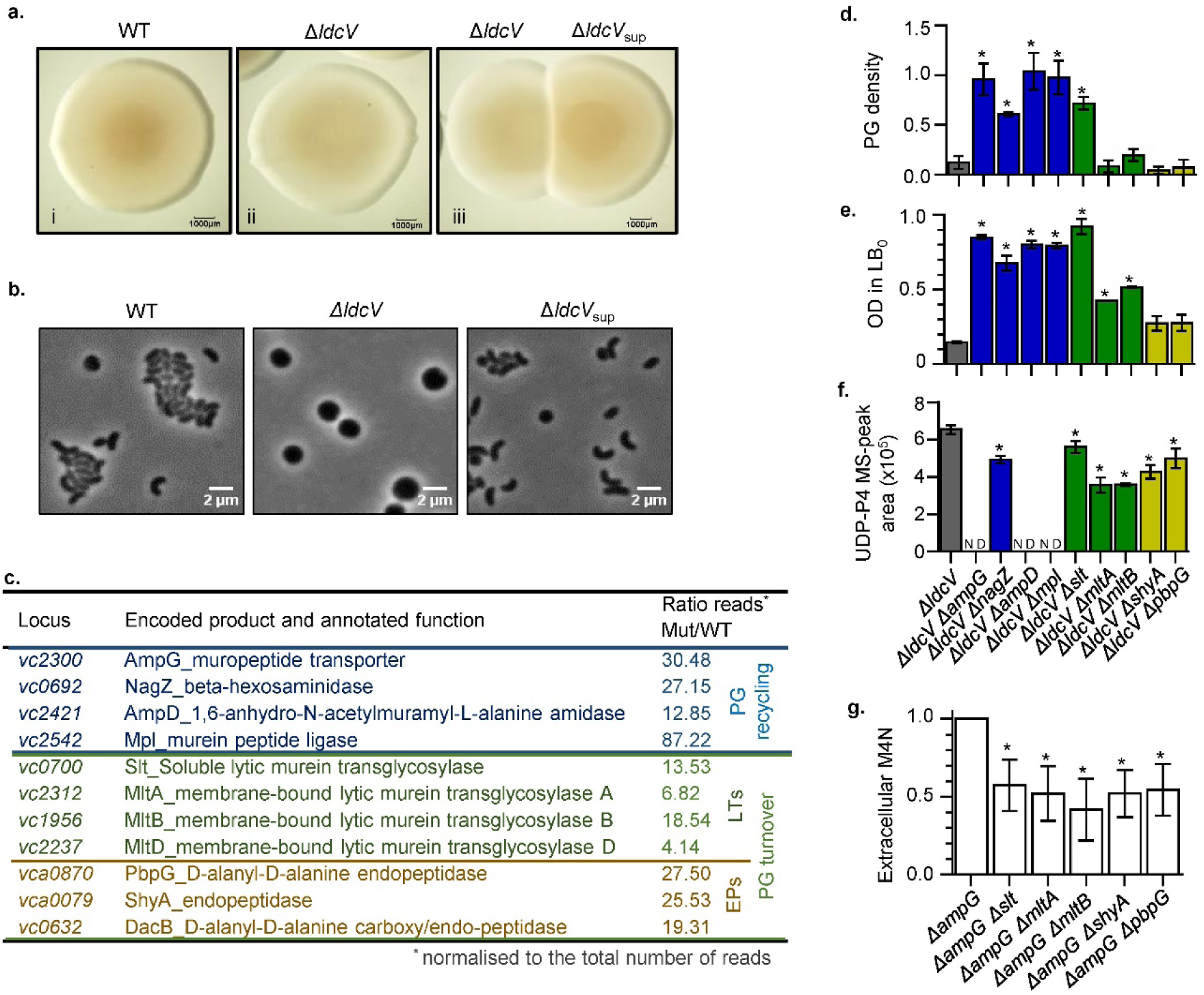
Aberrant phenotypes of the Δ*ldcV* mutant can be suppressed by inactivation of genes involved in PG-recycling or PG-turnover. **a-b.** Colony (**a**) and cell (**b**) morphology of wild-type, *ΔldcV* and a *ΔldcV* suppressor mutant (*ΔldcVsup*) growing in LB_10_**. c.** List of the suppressor candidate genes annotated to be involved in PG homeostasis obtained from sequencing the *ΔldcV-*LB_0_ Tn-library. **d-f.** Phenotypes of *ΔldcV* suppressor mutants: (**d**) amount of PG per cell (normalised to the WT levels) of stationary-LB_10_ cultures; (**e**) OD_600_ measured after 10h incubation in LB_0_; and (**f**) level of cytosolic UDP-P4 of the indicated mutant strains. g. Amount of soluble M4N (normalised to the M4N detected for the *ΔampG* single mutant) detected by MS^e^ in the extracellular medium of exponential cultures of indicated lytic transglycosylase or endopeptidase mutants. Data represent mean ± s.d. of 2 (e, g) or 3 (d, f) biological replicates; ND: not detected; \**P* < 0.05, unpaired t-test.

We also carried out a transposon insertion sequencing (Tn-seq) based screen to identify additional suppressors of *ΔldcV* lethality on LB0 agar plates. Notably, more than one third of the candidate suppressors corresponded to genes associated with PG homeostasis (**Supplementary Fig. 4a-b**). Insertions in genes associated with the PG recycling pathway^12^*, ampG, nagZ, ampD and mpl*, were identified as suppressors, consistent with the spontaneous suppressor data. In addition, this approach yielded Tn insertions in genes encoding diverse PG cleaving enzymes^8^ (**Fig. 2c, Supplementary Fig. 4c**).

Double deletion mutants, combining *ΔldcV* with deletions in most of identified suppressors, were constructed to validate and extend our analysis of the putative *ldcV* suppressors. Combining *ΔldcV* with individual deletions of PG recycling genes, *ampG* (permease)*, ampD* (amidase) or *mpl* (muropeptide ligase), yielded near complete reversion of the *ldcV* phenotypic defects, which correlated with the absence of detectable UDP-P4 (**Fig. 2d-f and Supplementary Fig. 5a-b**). However, the *ΔldcV ΔnagZ* mutant accumulated UDP-P4 and incompletely suppressed the growth and PG defects of the *ΔldcV* mutant (**Fig. 2d-f, Supplementary Fig. 5a-b**). UDP-P4 accumulation in the *nagZ ldcV* mutant is likely explained by AmpD’s catalytic promiscuity^30^, which can release the tetrapeptide from both the anhydro-muramyl-peptide and the M4N (accumulated in the *nagZ* mutant background) (**Supplementary Fig. 5c-d**) thereby promoting Mpl-dependent formation of UDP-P4. Similarly, deletions in PG-hydrolytic enzymes only partially alleviated the *ΔldcV* phenotypes and did not prevent UDP-P4 accumulation (**Fig. 2d-f**). Therefore, these data suggest that in addition to UDP-P4 depletion, there is a second mechanism to supress *ΔldcV* lethality.

PG hydrolases are also known as autolysins because their uncontrolled activity upon cessation of PG synthesis can lead to cell lysis^31,32,33^. Therefore, we hypothesized that inactivation of these enzymes could suppress *ΔldcV*’s viability defect by reducing autolysis. To monitor autolysis, we compared the amount of the PG turnover product M4N in the extracellular medium in the wild-type strain and the PG hydrolase mutants in a *ΔampG* background (to block PG recycling). There were reduced amounts of extracellular M4N in all PG hydrolase mutants (**Fig. 2g**), suggesting that the hydrolase deletion mutants may relieve the toxicity associated with the absence of LdcV by reducing PG turnover.

### Recycled tetrapeptides can be incorporated into peptidoglycan leading to reduced cross-linkage

Höltje and colleagues speculated that the lytic phenotype of an *E.coli* Δ*ldcA* mutant results from incorporation of UDP-P4 into PG, which in turn causes a lethal reduction in peptide cross-linkage^19^. However, incorporation of recycled tetrapeptides into the PG matrix has not been experimentally verified. Lipid II is the final precursor for PG synthesis^34^; notably, most Lipid II extracted from the *ldcV* mutant was in the tetrapeptide form rather than the native pentapeptide form found in the wild-type (**Fig. 3a, Supplementary Fig. 6**). To track whether recycled tetrapeptides can be incorporated into the mature PG sacculus, we designed a strategy to distinguish between tetrapeptides derived from the *de novo* pathway (ending with D-Ala) and those from the recycling pathway. To this end, the growth medium was supplemented with a “D-Met-labelled” anhydro-murotetrapeptide derivative (M4N^Met^) which can be recycled and detected as a D-Met tetrapeptide in the PG (**Fig. 3b, Supplementary Fig. 7a**). These experiments were carried out in a *ΔldcV* derivative (referred to as *Δ*4) that was also deficient in endogenous D-Met production (due to the absence of the BsrV racemase) and unable to incorporate exogenous D-Met by L,D-transpeptidation^4^ (due to the absence of LdtA and LdtB). M4^Met^ was detected in the PG isolated from the M4N^Met^-treated *Δ*4 strain, but not in a control strain (*Δ*5), where recycling was disabled via *ampG* inactivation (**Fig. 3c and Supplementary Fig 7b**). Together, these observations demonstrate that the recycled tetrapeptides that accumulate in the *ldcV* mutant are reused as substrates for PG synthesis.

**Fig. 3.**
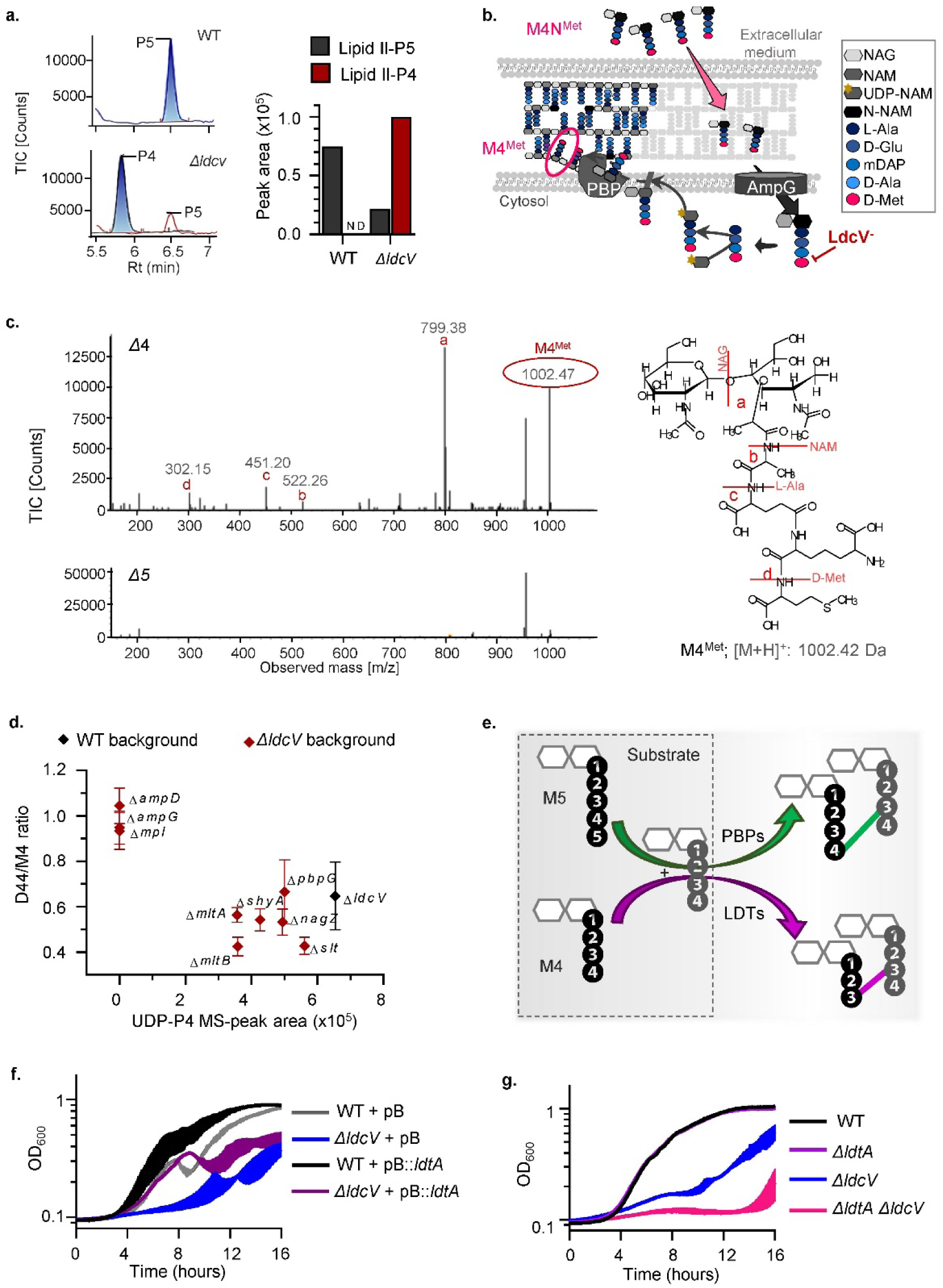
Incorporation of recycled UDP-P4 into the cell wall leads to a reduction in D,D-cross-linking. **a.** Detection and quantification of lipid-II P4 and P5 delipidated-forms isolated from WT and *ΔldcV* cultures. **b.** Schematic representation of the protocol used to demonstrate incorporation of recycled M4N^Met^ into peptidoglycan. **c**. Exogenous M4N^Met^ was added to cultures of *ΔldtA ΔldtB ΔbsrV ΔldcV* (*Δ*4) and *ΔldtA ΔldtB ΔbsrV ΔldcV ΔampG* (*Δ*5). The MS/MS profile obtained by targeted MS of PG derived from *Δ*4 but not *Δ*5 revealed the presence M4^Met^. **d.** Quantification of D,D-cross-linkage (estimated by calculating the D44/M4 ratio and normalised to the WT value) present in the PG of *ΔldcV* suppressors mutants (Y axis) and amount of accumulated UDP-P4 detected for those strains (X axis). **e.** Schematic representation of muropeptide cross-linkage mediated by penicillin binding proteins (PBPs) and LD-transpeptidases (LDTs). **f-g**. Growth kinetics in LB_0_ of *ΔldcV* mutant overexpressing *ldtA* from the arabinose-inducible P_BAD_ promoter in the pBAD_18_ vector (pB) (**f**) and of *ΔldtA ΔldcV* double mutant (**g**). ND: not detected.

In addition to accumulating tetrapeptide precursors, the *ΔldcV* mutants exhibited a marked reduction in D,D-cross-links (**Fig. 3d, Supplementary Fig. 8a**). We speculated that an increase in L,D-cross-links could compensate for the reduction in D,D-cross-links in the Δ*ldcV* mutant, since L,D-transpeptidases use tetrapeptides instead of pentapeptides as substrates^35, 36^ (**Fig. 3e**). Consistent with this idea, we found that expression of LdtA (the main L,D-transpeptidase of *V. cholerae*)^4^ increased the fitness of the *ΔldcV* mutant (**Fig. 3f, Supplementary Fig. 8b**), whereas deletion of *ldtA* in the *ΔldcV* background further attenuated its growth (**Fig. 3g**).

### Recycled tetrapeptides reduce the abundance of UDP-MurNAc-pentapeptide

Besides the reduction in PG cross-links, we noticed a striking correlation between accumulation of tetrapeptide precursors (Lipid II-P4 or UDP-P4) and a reduction in canonical pentapeptide precursors (**Fig. 3a and Fig. 4a**). This result is consistent with the reduced PG content of the *ldcV* mutant and suggests that recycled tetrapeptide precursors negatively regulate *de novo* PG biosynthesis. We hypothesized that the reduction of UDP-P5, derived from the *de novo* PG synthesis pathway, might be the result of UDP-muramyl (the shared intermediate) being consumed by the recycling pathway. Therefore, increasing the flow of the *de novo* pathway, i.e. the levels of UDP-P5 ^37^, should alleviate *ΔldcV*’s PG defect. Indeed, overexpression of any of the MurA-F enzymes increased the PG density of the *ldcV* mutant (2-4 times) and greatly improved its fitness (more than 1000 times in the case of MurC overexpression) in competition experiments (**Fig. 4b-c, Supplementary Fig. 8c**). Interestingly, among the Mur enzymes, expression of MurC had the most potent effect on improving *ldcV* mutant fitness. MurC mediates formation of UDP-muramyl-L-Ala, a substrate committed to UDP-P5 regardless of the presence of tetrapeptides. Therefore, our data suggest that competition between the *de novo* synthesis and recycling pathways for available UDP-muramyl accounts for the reduction in the amount of the UDP-P5 precursors in the ΔldcV mutant.

**Fig. 4.**
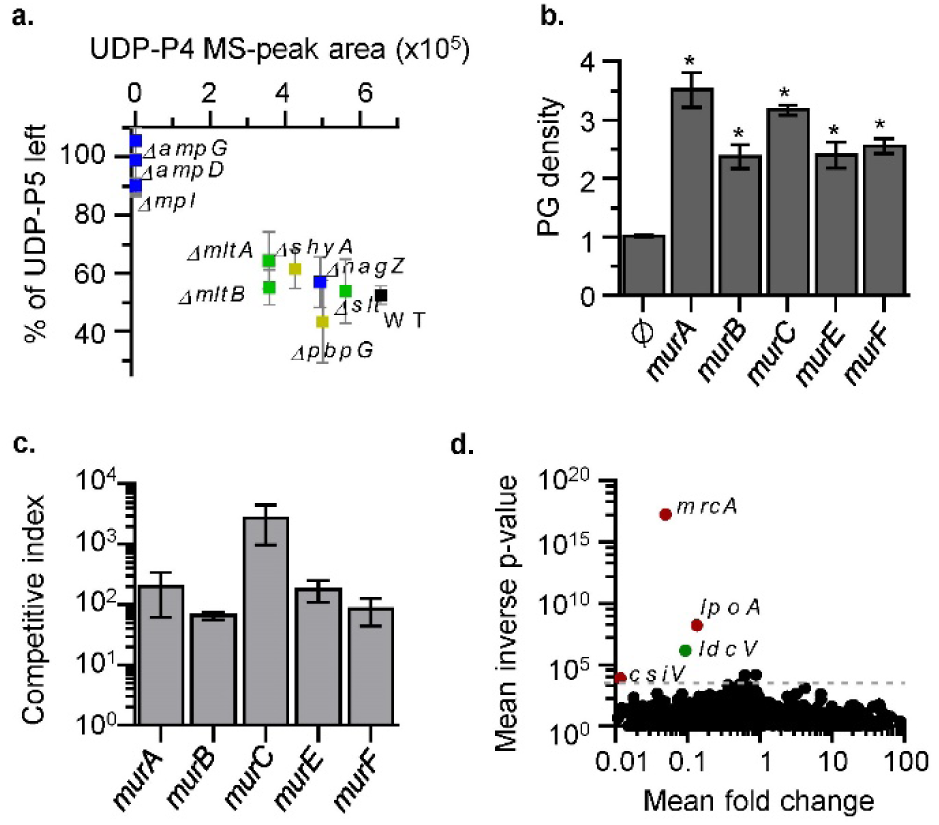
Negative interactions between the recycling and the *de novo* PG-biosynthetic pathways. **a.** Cytosolic UDP-P5 in indicated strains after deleting Δ*ldcV* with respect to the single mutant (Y axis) and detected cytosolic UDP-P4 after Δ*ldcV* deletion (X axis). **b.** PG density found in stationary LB_10_cultures of the *ΔldcV* mutant when overexpressing the indicated Mur locus from the pBAD_18_ vector (normalised to the value of the strain carrying the empty vector) (Ø: empty plasmid). **c.** Competition assay in LB_0_ of the *ΔldcV* mutant strains overexpressing the indicated Mur-proteins *vs.* the *ΔldcV* mutant carrying the pBAD_18_-empty vector. **d.** Volcano plot showing the ratio of read counts mapped to individual genes in the *ΔldcV-*mutant LB_10_-transposon library compared with the control wild-type LB_10_-library. Genes represented in red and green are significantly underrepresented. Data represent mean ± s.d. of 2 (c) or 3 (d) biological replicates; \**P* < 0.05, unpaired t-test.

Also consistent with the hypothesis that UDP-P4 negatively regulates pentapeptide levels, a Tn-seq based screen for *ldcV* genetic interactions revealed a synthetic lethal/sick phenotype with the genes encoding *V. cholera*e’s dominant high molecular weight PBP1A (*mrcA*) and its regulatory partner proteins LpoA and CsiV (*vc0581* and *vc1887* respectively) (Fig. 4d). Collectively, these observations buttress the idea that LdcV plays a key role in maintaining a proper ratio between UDP-P5 and UDP-P4 to ensure optimal PG synthesis and composition (i.e. cross-linking).

### Recycled non-canonical tetrapeptide precursors are detectable in wild-type *V. cholerae*

We previously found that PG editing by non-canonical D-amino acids (NCDAA) such as D-Met leads to accumulation of pentapeptides in the cell wall of some bacteria, likely because these modified muropeptides are poor substrates for D,D-carboxypeptidases^4^. We wondered whether NCDAA modified muropeptides might also be sub-optimal substrates for LcdV, and lead to accumulation of recycled tetrapeptides in the wild-type strain. Notably, targeted MS analysis of stationary phase wild-type cytosolic extracts revealed the presence of UDP-tetrapeptide with D-Met at the terminal position (UDP-P4^Met^) (**Fig. 5a and Supplementary Fig. 10**). In contrast, UDP-tetrapeptides were not detected in extracts from the *ΔampG* mutant, indicating that the source of these molecules is recycled anhydro-muropeptides (**Fig. 5a**). Furthermore, we directly detected the presence of anhydro-murotetrapeptides with D-Met (M4N^Met^) in the extracellular medium (**Fig. 5b**). Like UDP-P4^Met^, M4N^Met^ was exclusively detected in stationary phase (**Fig. 5b and Supplementary Fig. 9a**), consistent with the stationary phase–limited expression of BsrV, the racemase that generates D-Met^4^. Similarly, M4N^Met^ was not detected in supernatants from a *ΔbsrV* strain (**Supplementary Fig. 9b**).

**Fig. 5.**
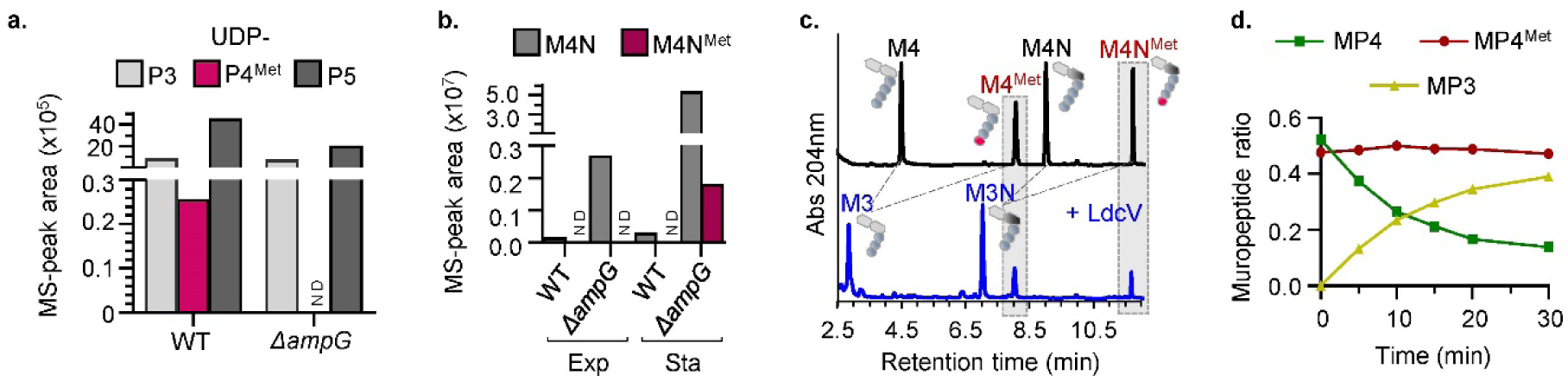
LdcV exhibits a preference for hydrolysing canonical over D-Met modified murotetrapeptides leading to accumulation UDP-P4 precursors. **a.** UDP-forms detected by targeted MS in the cytosolic extract of stationary phase cultures of indicated strains growing in LB+L-Met. **b.** Extracellular anhydro-murotetrapeptides detected by targeted MS in exponential (Exp) or stationary (Sta) cultures of indicated strains growing in LB+L-Met. **c**. UPLC chromatograms showing the substrate (in black) and the resultant products (in blue) of LdcV *in vitro* assays (1h incubation) (M4: murotetrapeptide; M4^Met^: murotetrapeptide with D-Met at fourth position; M4N: anhydro-murotetrapeptide; M4N^Met^: anhydro-murotetrapeptide with D-Met at fourth position; M3: murotripeptide; M3N: anhydro-murotripeptide). **d.** Dynamics of substrate consumption during *in vitro* reactions of LdcV (MP4= M4 + M4N; MP4^Met^= M4^Met^ + M4N^Met^; MP3= M3 + M3N). ND: not detected.

To assess if accumulation of UDP-P4^Met^ could be due to LdcV’s preference for canonical vs. non canonical tetrapeptides (D-Met modified), purified LdcV was incubated with a mixture of canonical (D-Ala at fourth position, M4N and M4) and non-canonical (D-Met at fourth position, M4N^Met^ and M4^Met^) murotetrapeptides as potential substrates. Analyses of the digestion products revealed that while LdcV can hydrolyse both kind of substrates (**Fig. 5c**), it exhibits a preference for canonical over D-Met-modified tetrapeptides (**Fig. 5d, Supplementary Fig. 9c**). The preference of LdcV for canonical tetrapeptide substrates, could lead to accumulation of non-canonical tetrapeptide precursors in the wild-type strain (functional LdcV). Furthermore, additional LdcV-like (from *Aeromonas hydrophila* and *Proteus mirabilis*) and LdcA-like (from *E. coli* and *Salmonella enterica*) L,D-carboxypeptidases exhibit similar preference for canonical rather than NCDAA-modified murotetrapeptides substrates (**Supplementary Fig. 9d**). Thus, the recycling of NCDAA-edited PG and subsequent production of precursor tetrapeptides could underlie a conserved mechanism linking NCDAA editing with the control of PG synthesis.

## DISCUSSION

The general biological importance of the PG recycling pathway is not well understood, since its inactivation has little impact on bacterial growth under laboratory conditions^11, 38^. Here, our elucidation of functions of LdcV, *V. cholerae’s* L,D-carboxypeptidase, suggests that recycling and L,D-carboxypeptidases in particular exert a critical role in PG homeostasis, modulating PG synthesis and composition.

A *V. cholerae ldcV* mutant exhibited reduced PG content, spherical morphology and sensitivity to low osmolarity. Assays with purified LdcV established that the protein has L,D-carboxypeptidase activity, although it was annotated as a D,D-carboxypeptidase and lacks homology to *E. coli’*s LdcA. Analyses of suppressors of the *ldcV* mutant’s phenotype revealed that *ldcV* plays an important role in the gene network controlling PG homeostasis. There appears to be at least 2 related pathways – PG recycling and PG cleavage – through which suppressor mutants mediate their effects. Some suppressors (e.g., inactivation of *ampG* or *mpl*) prevented UDP-P4 formation, thus preventing its toxic effects on PG biogenesis, whereas others (e.g. inactivation of *slt*) reduced PG turnover. Moreover, our observations suggest that the lethality of Δ*ldcV* is caused by two distinct mechanisms: 1) incorporation of recycled UDP-P4 into the murein (**Fig. 3**), which leads to a decrease in PG cross-linkage; and 2) a reduction of the canonical UDP-P5 precursor pool (**Fig. 4**), which leads to a reduction of PG synthesis.

Incorporation of recycled tetrapeptides into the murein sacculus has not been demonstrated previously. Distinguishing tetrapeptides that are generated from the activities of periplasmic PBP transpeptidases/carboxypeptidases on pentapeptide substrates from those derived from flipped lipid II tetrapeptides was challenging. Provision of exogenous traceable D-Met anhydromuropeptides derivatives to a *ΔldcV* mutant strain incapable of synthesizing NCDAA-modified muropeptides, enabled detection of recycled tetrapeptides in the PG. The use of non-canonical anhydromurotetrapeptides (e.g. fluorescent derivatives) could facilitate the screening of multiple species for the absence/presence of PG recycling pathways and also to assess Ldc substrate specificity *in vivo*. Furthermore, our findings suggest that using compounds that resemble NCDAA-modified tetrapeptides could be a means to inhibit PG biogenesis; notably, the D-Ser residue in the β-lactam antibiotic nocardicin A is thought to specifically target Lcds^39^.

Lipid-II tetrapeptide substrates can only serve as acceptors in PBP-mediated transpeptidase reactions and thus their incorporation into the sacculus results in reduced PG cross-linkage. Consistent with incorporation of substrates unsuitable for the high molecular weight PBPs, we found that LdtA, an L,D-transpeptidase that uses tetrapeptides as donor substrates^4^, can compensate for the growth defect of the *ldcV* mutant in low osmolarity (**Fig. 3, Supplementary Fig. 8b**). Interestingly, a number of bacterial species lack Ldc orthologues, despite apparently encoding other components of the PG recycling pathway (e.g. *Yersinia pestis* and *Acinetobacter baumanni*^12^). In organism lacking Ldc homologues, we hypothesize that L,D-transpeptidases (Ldts) may play more prominent roles in maintaining PG homeostasis. Moreover, our data suggests that inhibition of Ldc (e.g., with nocardicin A) in combination with inhibitors of Ldts (e.g., imipenem)^40^ and/or PBP1a (e.g. beta-lactams) might have synergistic effects and thus could represent a potent drug combination for antimicrobial therapy.

Cytosolic accumulation of recycled tetrapeptides is largely prevented by the efficient cleavage activity of LdcV. However, tetrapeptides can accumulate when they are modified by NCDAAs (e.g. D-Met), since they are less preferred substrates for L,D-carboxypeptidation compared to their canonical counterparts (ending with D-Ala). Indeed, we detected accumulation of UDP-tetrapeptide precursors modified with D-Met (UDP-P4^Met^) in stationary phase wild-type cultures (**Fig. 5**), demonstrating that in *V. cholerae* stationary phase production of D-Met fine-tunes PG synthesis activity via the recycling pathway. Thus, production of NCDAA modulates cell wall synthesis and cross-linking by at least two mechanisms – 1) through periplasmic PG editing (by Ldts) with free NCDAA^3–5^; and 2) incorporation of recycled NCDAA-modified tetrapeptide precursors – that together coordinate cell wall synthesis with growth arrest (**Fig. 6i**). It is also possible that in addition to auto-regulatory roles, production of NCDAA induces regulatory changes in the murein of neighbouring organisms (**Fig. 6ii-iii**). In principle, any bacterium that incorporates NCDAA in the 4^th^ position of the muropeptides (e.g. via the activity of Ldts) is a potential source of NCDAA-modified anhydromuropeptides that could modulate PG synthesis/composition in bacteria inhabiting the same niche (**Fig. 6**).

**Fig. 6.**
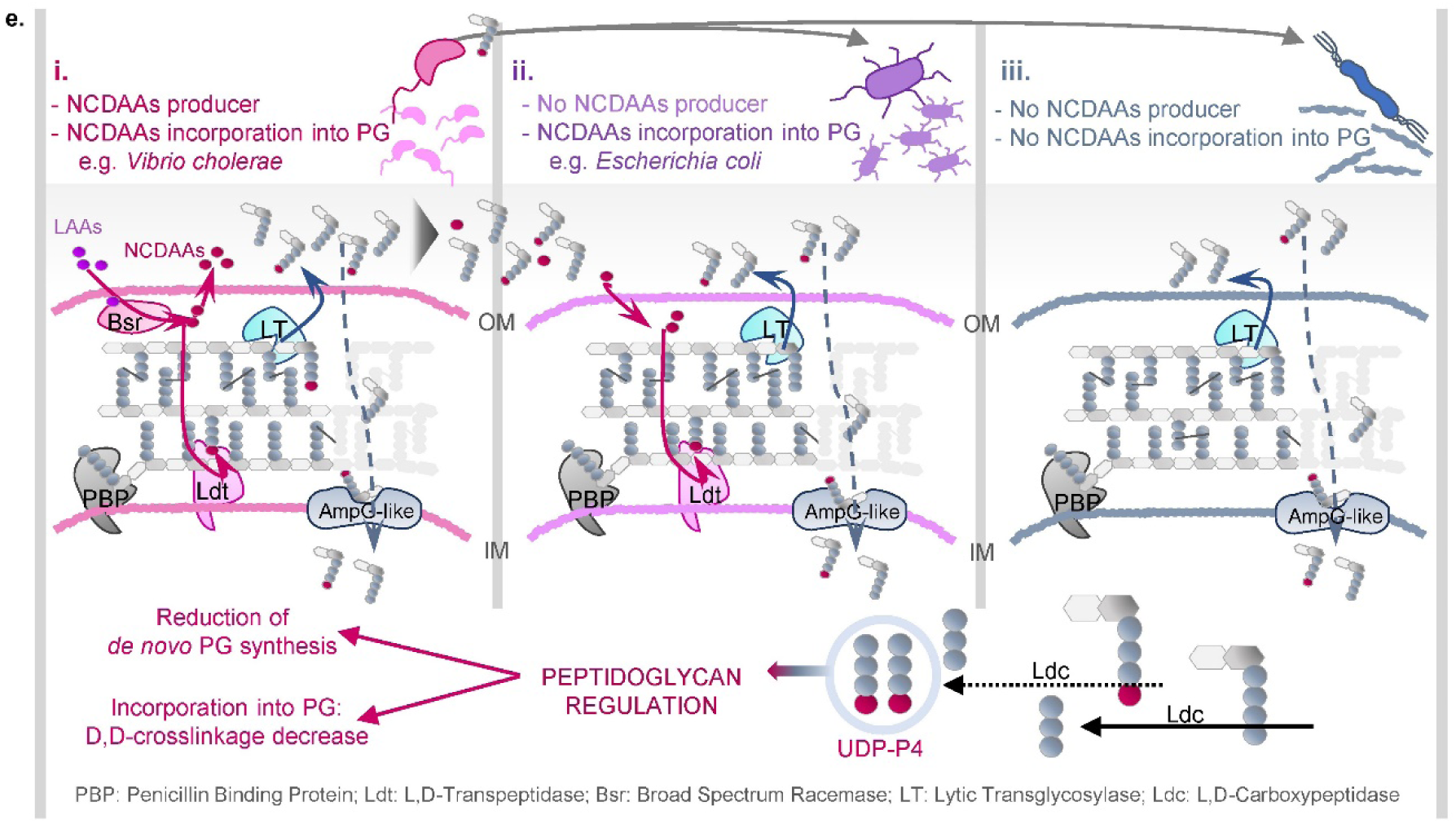
Model of regulation of PG-synthesis by physiological production of UDP-P4 in a multispecies bacterial environment.

Finally, extracellular PG fragments are known to be important signals in innate immunity, organ development and behaviour^41–45^. Our observation that bacteria can release PG fragments modified with NCDAA suggests that it will be important to consider whether NCDAA-modified PG fragments convey distinct information in inter-kingdom signalling compared to fragments modified with canonical DAA. Moreover, in microbial ecology (**Fig 6**), our findings suggest that release of extracellular non-canonical muropeptides could mediate interspecies modulation of PG synthesis in ‘trans’ if such peptides become substrates for PG recycling in neighboring organisms.

## METHODS

### Bacterial strains, growth conditions, and construction of mutants and plasmids

Strains are listed in **Supplementary Table 1**. All *V. cholerae* strains used in this study are derivatives of the sequenced El Tor clinical isolate N16961^46^. *E. coli* strains DH5α and DH5α λ*pir* were used as hosts for constructing plasmids. Unless otherwise specified, bacteria were grown at 37 °C in LB (Luria Bertani broth) medium. Agar 1.5% (w/v) was used in solid plates. Antibiotics were used at the following concentrations (per ml): streptomycin (Sm), 200 μg, ampicillin (Ap), 50 μg, carbenicillin (Cb), 50 μg, chloramphenicol (Cm), 20 μg (*E. coli*) and 5 μg (*V. cholerae*), and kanamycin (Kn), 50 μg. L- or D-methionine (L/D-Met) were used at a final concentration of 20 mM.

*V. cholerae* deletion mutant strains were constructed using standard allele exchange techniques with derivatives of the suicide plasmid pCVD442 as described previously^47^. Primers used for constructing pCVD442-derivatives (primers P1-4) and for verifying the deletions (external primers E1-2) are shown in **Supplementary Table 2**. Overlaping extension PCR was carried out as described^48^. Since the start of *vc2153* overlaps with the terminal 73 nucleotides of the essential locus *vc2152*^49^, the *Δvc2153* mutant removes *vc2153* nucleotides 171 to 747. Because strains bearing *vc2153* mutations accumulate suppressors, the deletion of *vc2153* was always the final mutation introduced in strains bearing inactivation of more than one gene.

Complementation and overexpression plasmids were constructed using the primers indicated in **Supplementary Table 3** by amplifying the gene of interest with its native ribosome binding site. The *ldcA* gene from pET28b*::ldcA* was inserted into pBAD33 yielding pBAD33*::ldcA.* Expression of cloned loci from the PBAD promoter was induced by addition of 0.2% arabinose^50^.

For growth curves, stationary cultures were normalized to an optical density at 600 nm (OD600) of 1.5, diluted 1:100 and used for inoculating 96-well plates containing 180 μl of fresh medium. At least three replicates per strain and condition were carried out. Measures of the OD600 for growth curves were carried at 30°C with shaking using an Eon Biotek microplate spectrophotometer.

DNA and protein sequence alignments were carried out using the blastn and blastp programs (http://www.ncbi.nlm.nih.gov/BLAST/), respectively.

### Cell and colony morphology imaging

Analysis of cell morphology was performed on immobilized bacteria (1% agarose LB pads) by phase-contrast microscopy using a Zeiss Axio Imager.Z2 microscope equipped with a 100x oil immersion objective and a Hamamatsu digital camera controlled by Zeiss Zen Blue software. Pictures of colonies were taken from LB10 plates (10 g L^−1^ of NaCl) by using a Nikon SMZ1500 Zoom Stereomicroscope and a DS-Fi1 High-Definition Color Camera. Brightness and contrast levels of the images were adjusted using ImageJ software.

### Isolation and analysis of peptidoglycan amount and composition

Murein sacculi isolation and muropeptide analysis were performed essentially as described previously^51–53^. Briefly, bacterial pellets of 50 ml (exponential phase OD600 ∼0.4) or 25ml (stationary phase OD600 ∼3) cultures were boiled in 5% SDS. Cell wall material was pelleted and repeatedly washed with water by ultracentrifugation. Clean sacculi were digested with muramidase (100 µg/ml) and soluble muropeptides were then reduced using 0.5 M sodium borate pH 9.5 and sodium borohydride at 10 mg ml^−1^ (final concentration). The pH of the samples was adjusted to 3.5 with phosphoric acid for liquid chromatography.

UPLC analyses were performed on a Waters-UPLC system equipped with an ACQUITY UPLC BEH C18 Column, 130Å, 1.7 µm, 2.1 mm×150 mm (Water, USA) and detected at Abs. 204 nm. Muropeptides were separated primarily using a linear gradient from buffer A (phosphate buffer 50 mM pH 4.35) to buffer B (phosphate buffer 50 mM pH 4.95 methanol 15% (v/v)) in a 20 min run; a modified method using organic solvents (described below in the LC-MS analysis methods) was used for analysis D-Met muropeptides for a better separation of the peaks and to avoid co-elution of muropeptides. Identity of the peaks was assigned by comparison of the retention times and profiles to other chromatograms in which mass spectrometry data had been collected, and by mass spectrometry when necessary. The relative amount of each muropeptide was calculated by dividing the peak-area of a muropeptide by the total area of the chromatogram. The apparent density of PG was assessed by normalizing the total area of the chromatogram to the OD600 of the culture used for the PG purification. The degree of cross-linking was calculated as described previously^52^ and as a rough estimation of D,D-cross-linkage the ratio between the tetra-tetra dimer (D44) and the tetra monomer (M4) was calculated. All PG analyses were preformed using biological triplicates.

### Protein overexpression and purification

*vc2153* and the genes encoding for the cytoplasmic L,D-carboxypeptidases of *E. coli* (LdcA), *A. hydrophila* (AHA_1477) *P. mirabilis* (PMI1557) and *S. enterica* (STM1800) were cloned into pET28b and *vca0337* into pET22b (Novagen) for expression in *E.coli* BL21(DE3) cells (primers used for cloning are listed in **Supplementary Table 4**). Expression was induced in LB cultures at exponential phase with 1 mM isopropyl β-D-thiogalactopyranoside (IPTG) for 2 h. Cells were harvested, resuspended in 150 mM Tris HCl pH 7.5, 150 mM NaCl, and stored at −20 °C. After thawing on ice, cells were disrupted by passaging through a French press twice. 6×His-tagged proteins were purified from cleared lysates (30 min, 50,000 rpm) on nickel-nitrilotriacetic acid-agarose columns (Qiagen), and eluted with 150 mM Tris HCl pH 7.5, 150 mM NaCl, 250 mM imidazole. The eluate was dialyzed for 12 h in 50 mM Tris-HCl pH 7.5, 100 mM NaCl. Purified proteins were visualized by SDS-PAGE and Coomassie Brilliant Blue staining and quantified by Bio-Rad Protein Assay (Bio-Rad).

### Substrate preparation and *in vitro* reactions for protein activity assays

Isolated *V. cholerae* or *E. coli* PG was digested with muramidase (see method above) and used for purification of muropeptides by collection of HPLC-separated muropeptide peaks. For isolation of anhydro- or D-Met-modified muropeptides, *in vitro* reactions were performed using purified Slt70 and LdtA proteins respectively^4, 54^. Acetonitrile and formic acid were used as organic solvents for peak separation. Collected peaks were lyophilized, dissolved in water, and stored at −20 °C.

*In vitro* reactions were prepared in a final volume of 50μl containing as substrates either pure muropeptides (10ug), intact sacculi or muramidase-digested sacculi; 10ug of purified protein (50 mM final concentration) in Tris-HCl pH 8 buffer was used. Reactions were performed for 60 min at 37°C, heat inactivated (100 °C, 15 min) and centrifuged (15,000 rpm, 10 min) to remove precipitated material. Peptides in the supernatant were monitored by UPLC, identified using standard controls based on their retention time, and confirmed by UPLC-MS analysis.

For studing the substrate preference of Ldcs for different muropeptides, M4, M4^Met^, M4N and M4N^Met^ were produced and purified as described above. Then a mixture of the four muropeptides (containing approximately equal amounts of each of them) was used as substrate in a reaction of 150 ul final volume in 50mM Tris-HCl pH 8 buffer and 50 ug of purified Ldc protein. After addition of the protein, samples were incubated at 37° C and small aliquotes were removed at several incubation times, boiled to inactivate the protein and centrifuged. Soluble products were injected into the UPLC. The ratio of each muropeptide at each time-point was calculated by dividing its area by the total area calculated for all the muropeptides (substrates and products).

### Whole genome sequencing of spontaneous suppressor mutants

Genomic DNA was extracted using the GeneJET Genomic DNA Purification Kit and quantified with Qubit dsDNA HS Assay Kit (Thermo Scientific). 1 ng of each DNA was used to generate the genomic libraries following the manufacturer’s recommendations (Nextera XT DNA Sample Preparation Kit, Illumina). DNA libraries were then pooled in equimolar proportions and sequenced employing a MiSeq Reagent Kit V2 (Illumina). Paired-end 2×300 bp reads were generated on an Illumina MiSeq instrument. Sequences of the isogenic wild-type strain N16961 and the *Δvc2153* mutant were determined in parallel.

The sequences were analyzed using the open, web-based computational platform Galaxy (https://usegalaxy.org/)^55^ as indicated in Alvarez *et al.*^3^. Mapping trimmed reads to the reference *V. cholerae* N16961 genome was performed by the BWA-MEM^56^ algorithm. After read alignment, Picard tools were used for the parent strains and each suppressor to mark and remove duplicate sequences mapping to different regions. Subsequently, SNPs (single-nucleotide polymorphisms) and indels (insertions and deletions) were detected by comparison of the results obtained after detecting genetic variants by FreeBayes^57^ and VarScan^58^. Mapping of the larger deletion in the suppressor mutant Sup4 was carried out using the Integrative Genome Viewer (IGV)^59^.

### Transposon insertion sequencing

*V. cholerae* transposon insertion libraries, of ∼150,000 insertion mutants each, were generated in N16961 and in the *Δvc2153* mutant as described previously^60^ using the Himar delivery vector pSC189^61^. Transposon mutants were plated directly onto LB10 or LB0 agar plates containing Sm and Km. Transposon insertion sequencing was performed as described previously^49^ using an Illumina MiSeq benchtop sequencer. Data analysis was conducted as described previously^49, 62^. Visual inspection of transposon insertion profiles was performed with the Sanger Artemis Genome Browser and Annotation tool^63^.

### Collection of extracellular and intracellular pools of soluble-muropeptides and LC-MS analysis

Sample preparation to determine the level of the different soluble muropeptides was performed following the protocol described previously by Lee *et* al.^64^ with some modifications. Briefly, bacteria were grown until exponential phase (OD600 ∼0.7), cooled on ice for 10 min and then, after adjusting the OD600 of the cultures (to have the same number of bacteria in each sample), normalized volumes of cells were harvested by centrifugation at 4,000 rpm, 4°C for 20 min. For analysis of the extracellular soluble muropeptides (ESM) normalized volumes of supernatants were collected, boiled for 15 min, centrifuged to remove precipitated material and stored at −20 °C. The cell pellets were gently resuspended and washed with ice-cold 0.9% NaCl solution. After pelleting the cells again by centrifugation, they were resuspended in the remaining volume and boiled in water for 15 min. Samples were centrifuged to remove cell debris at 14,000 rpm for 15 min, and soluble fractions (containing intracellular soluble muropeptides, ISM) were transferred to new tubes and stored at −20 °C. Both, ESM and ISM samples were filtered with 0.2 μm pore size filters, dried by speed vacuum, resuspended in water and used for LC-MS analyses. Soluble muropeptide analyses were performed on biological triplicates.

Detection and characterization of soluble muropeptides by LC-MS was performed on an UPLC system interfaced with a Xevo G2/XS Q-TOF mass spectrometer (Waters Corp.). Chromatographic separation was achieved as described previously^53^ using an ACQUITY UPLC BEH C18 Column (Waters Corp.) heated at 45 °C. 0.1% formic acid in Mili-Q water (Buffer A) 0.1% formic acid in acetonitrile (buffer B) were used as eluents. The gradient of buffer B was set as follows: 0-3 min 5%, 3-6 min 5-6.8%, 6-7.5 min 6.8-9%, 7.5-9 min 9-14%, 9-11 min 14-20%, 11-12 min hold at 20% with a flow rate of 0.175 ml/min; 12-12.10 min 20-90%, 12.1-13.5 min hold at 90%, 13.5-13.6 min 90-2%, 13.6-16 min hold at 2% with a flow rate of 0.3 ml/min; and then 16-18 min hold at 2% with a flow rate of 0.25 ml/min. Chromatograms were recorded at 204 nm. The QTOF-MS instrument was operated in positive ionization mode. Detection of ISM and ESM was in general performed by MS^e^ to allow the acquisition of precursor and product ion data simultaneously without pre-selection of targeted molecules. For MS^e^ the following parameters were set for ESI: capillary voltage at 3.0 kV, source temperature to 120 °C, desolvation temperature to 350 °C, sample cone voltage to 40 V, cone gas flow 100 L h^−1^ and desolvation gas flow 500 L h^−1^. Detection of D-Met modified tetrapeptides was achieved by targeted MS, by setting the collision energy to scan between 6 eV and 15–40 eV. Mass spectra were acquired at a speed of 0.25 s/scan. The scan was in a range of 100–2000 m/z. Data acquisition and processing was performed using UNIFI software package (Waters Corp.).

The molecular structure of each anhydro-muropeptide and the PG precursor molecules was obtained using ChemSketch^65^ to build a compound library in UNIFI. This compound library was used for processing the data, to detect and identify each molecule. Subsequent identification and confirmation of each muropeptide was performed by comparison of the retention-times and mass spectrometric data of experimental samples to purified authentic standards when available. Quantification was done by integrating peak areas from extracted ion chromatograms (EICs) of the corresponding m/z value of each muropeptide.

### Lipid II extraction and preparation for LC-MS analysis

Lipid extraction was performed according to the protocol described by Quiao *et* al.^66^ for “large-scale lipid extraction” but using only 500 ml cultures. Strains were grown in LB at 37 °C until an OD600 of 0.4-0.6 or 2 for exponential or stationary phase samples respectively. After the chloroform/methanol extraction steps, dried recovered interface fractions were resuspended in DMSO. The lipid tail of the extracted Lipid II was removed by ammonium acetate treatment before LC-MS analysis that was performed by targeted MS using the same parameters indicated above for analysing soluble-muropeptides pools.

### M4^Met^ incorporation into the PG by recycling

M4N^Met^ muropeptides were generated by initially obtaining M4N from SDS-free *V. cholerae* sacculi digested with purified *E. coli* His6-tagged Slt70 protein^54, 67^ and then replacing the terminal D-Ala by D-Met via *in vitro* L,D-transpeptidation mediated by purified LdtA protein in the presence of 20 mM D-Met^4^. Around 15 ug of M4N^Met^ (in water) or the same volume of water as negative control were added to 200ul of concentrated exponential cultures (containing ∼10^9^ bacteria) of *ΔldtA ΔldtB ΔbsrV ΔldcV* (Δ4) and *ΔldtA ΔldtB ΔbsrV ΔampG ΔldcV* (Δ5) strains grown in M9 minimal medium with glucose as a carbon source. After incubation of the samples for 45 min at 37°C, bacteria were pelleted and boiled in 2.5% of SDS for sacculi isolation and muramidase digestion. Solubilized muropeptides were reduced and after adjusting the pH, injected into the UPLC/MS for the search of M4^Met^ muropeptides by targeted MS.

### Competition assay of *ΔldcV* strains overexpressing Mur proteins

*In vitro* competition indices (CI) were determined from cultures containing the *ΔldcV* strain (*lacZ^+^*) carrying the empty pBAD18 vector and the *ΔldcV* (*lacZ^−^*) with the specified pBAD18::Mur-construction, mixed in a ratio 5:1. The competition assays were performed by incubating the cultures in LB0 and appropriate antibiotics supplemented with 0.2% of arabinose for 4h at 37 °C. After this time, CFUs were counted by plating serial log dilutions on LB0 agar containing Sm, Km, 0.2% arabinose and X-gal 40 μg ml^−1^. The CI was defined as the number of white colonies (strains overexpressing Mur proteins)/number of blue colonies (empty vector) counted after the incubation, divided by the white/blue ratio measure in the inoculum. The competition assays were performed in triplicates.

### Data analysis

GraphPad Prism 6 software was used for graphing data and statistical analysis.

## ACKNOWLEDGEMENTS

We thank all the members of the Cava lab for helpful discussions, especially Akhilesh K. Yadav and Akbar Espaillat for the support with the biochemistry part and Laura Alvarez for helping in the WGS and Tn-seq analysis. We also thank Veerasak Srisuknimit for his helpful comments on the manuscript.

Research in the Cava lab is supported by the by The Swedish Research Council (VR), The Knut and Alice Wallenberg Foundation (KAW), The Laboratory of Molecular Infection Medicine Sweden (MIMS) and The Kempe Foundation.

S.B.H. was supported by a Martin Escudero Postdoctoral fellowship.

Research in the Dörr lab is supported by National Institutes of Health (NIH) grants R01AI143704 and R01GM130971.

Research in the Waldor lab is supported by NIH grant RO1AI-042347 and HHMI.

## AUTHOR CONTRIBUTION

S.B.H. and F.C. designed and performed research and analysed data. F.C., S.B.H., T.D. and M.K.W. wrote the paper.

