## Supplementary Information for "Modulation of peptidoglycan synthesis by recycled cell wall tetrapeptides"

**This document includes:**

Supplementary Figures 1-10

Supplementary Tables 1-4

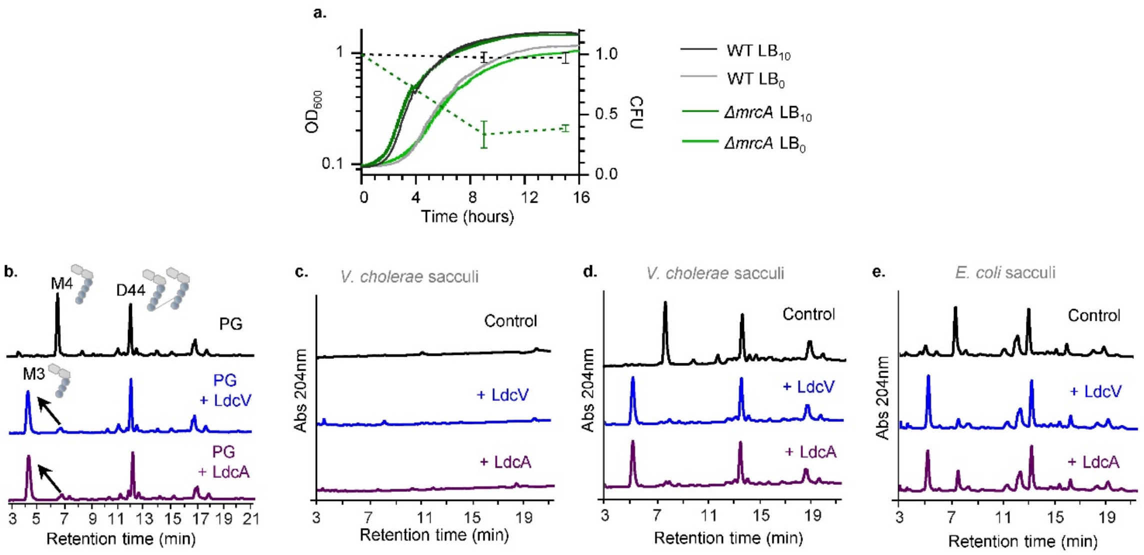


f.

| Identity Matrix (%) | LdcV | LdcA | Vca0337 |
|---|---|---|---|
| LdcV | 100.0 | 18.43 | 14.75 |
| LdcA | 18.43 | 100.00 | 30.42 |
| Vca0337 | 14.75 | 30.42 | 100.00 |

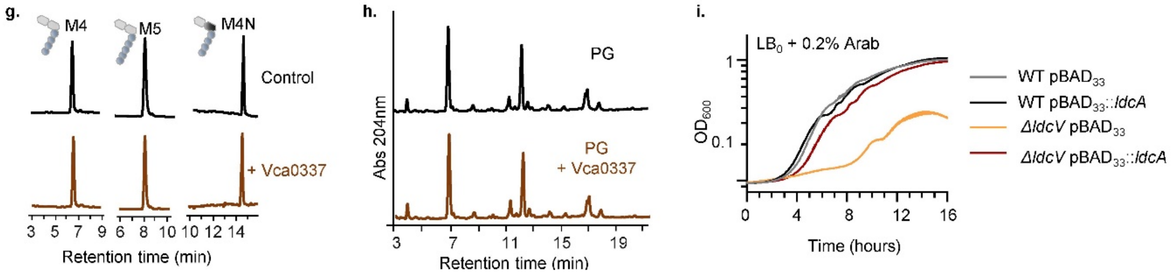


**Supplementary Fig. 1. Vc2153 is the functional homologue of *E.coli* L,D-carboxypeptidase LdcA. a.** Growth kinetics of indicated strains growing in LB containing 0 ($LB_0$) or 10 ($LB_{10}$) g/L of NaCl (left Y axis) and amount of CFU (normalised to wt) counted from $LB_{10}$ cultures at indicated times (dashed lines, right Y axis). **b.** UPLC chromatograms showing the products resulting after 90 minutes incubation of purified LcdV and LdcA with muramidase-digested PG. **c-e**. UCPL chromatograms showing soluble muropeptides resulting after incubation of indicated purified proteins with isolated high molecular mass murein sacculi (**c**) and after digesting the non-solubilized product with muramidase (**d-e**). **f.** Percentage of amino acid sequence-identity between indicated proteins obtained by Clustal alignment. **g-h.** Results of assaying the *in vitro* activity of Vca0337 protein using as substrate different isolated monomeric muropeptides (M4: murotetrapeptide; M5: muropentapeptide; N: anhydro form of the muropeptide) (**g**) or with muramidase-digested PG (**h**). **i.** Cross-complementation of *ΔldcV* by the expression of *E. coli* locus *ldcA* from the $P_{ARA}$ promoter of $pBAD_{33}$ vector.

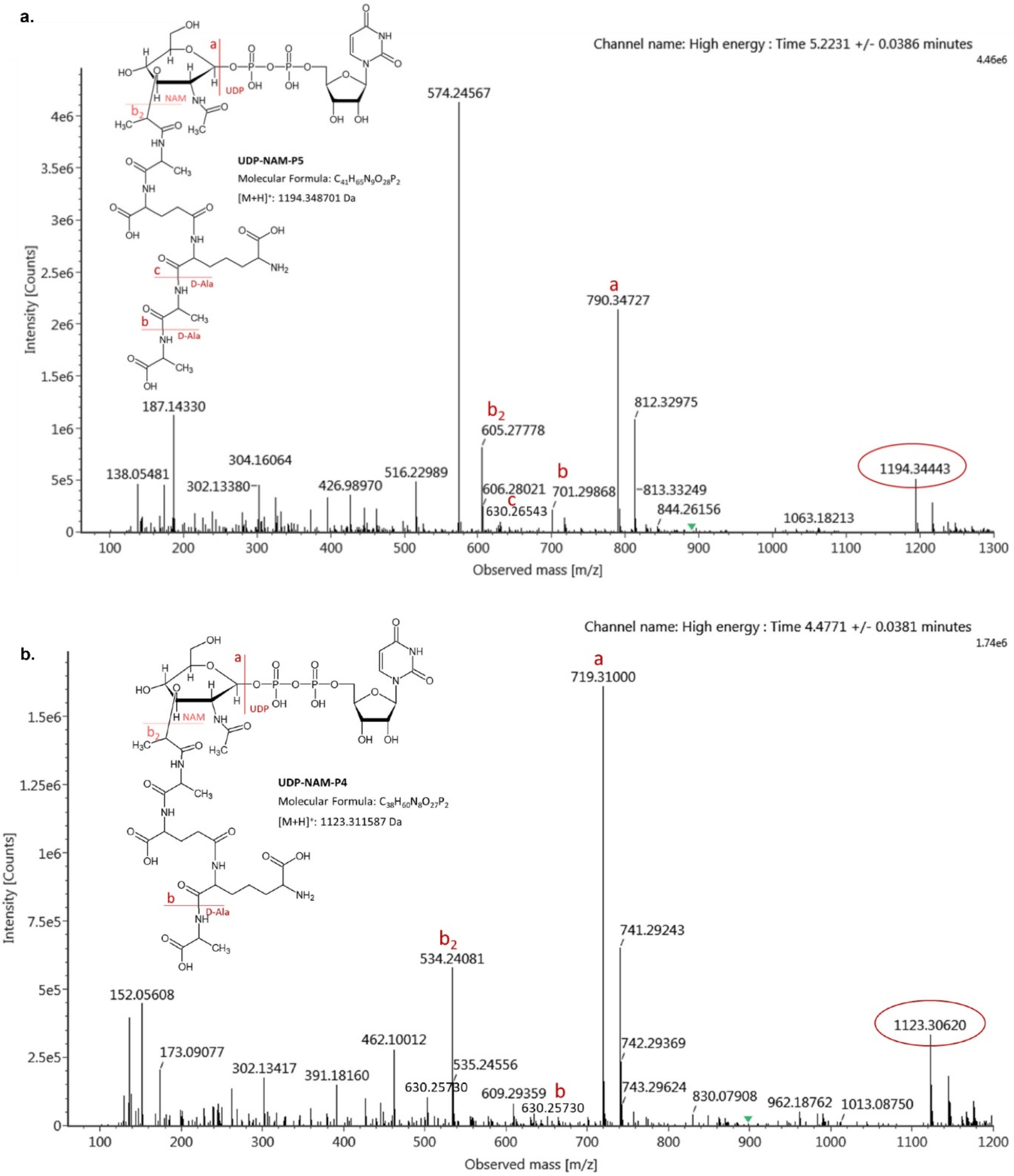


**Supplementary Fig. 2. Detection of UDP-tetrapeptide precursor accumulated in *ΔldcV* mutant.** MS/MS profile of UDP-P5 (**a**) and UDP-P4 (**b**) detected by untargeted MS in the cytosolic pool of PG-precursors of the WT (**a**) or the *ΔldcV* mutant (**b**) growing in LB.

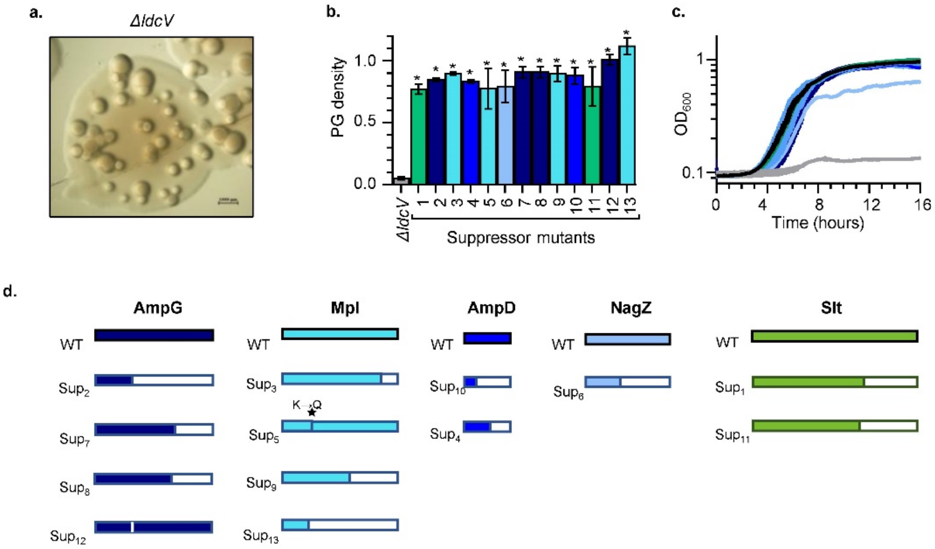


e.

| Strain | Mutation position[a] | Locus | Encoded protein | Nucleotide change[b] | Protein change[b] |
|---|---|---|---|---|---|
| **Sup$_1$** | Chr1_749695 | *vc0700* | Soluble lytic transglycosylase | G > A | Premature stop codon |
| **Sup$_2$** | Chr1_2454496 | *vc2300* | Permease AmpG | CGG > CG | Premature stop codon |
| **Sup$_3$** | Chr1_2723514 | *vc2542* | Murein peptide ligase Mpl | GAGCGCGGCTCAA > GA | Premature stop codon |
| **Sup$_4$** | Chr1_2595797 | *vc2421* | Amidase AmpD | Δ269-363 | Premature stop codon |
| **Sup$_5$** | Chr1_2724256 | *vc2542* | Murein peptide ligase Mpl | T > C | $K_{114}$→$Q_{114}$ |
| **Sup$_6$** | Chr1_740273 | *vc0692* | Glucosaminidase NagZ | G > T | Premature stop codon |
| **Sup$_7$** | Chr1_2455014 | *vc2300* | Permease AmpG | ATT > AT | Premature stop codon |
| **Sup$_8$** | Chr1_2455004 | *vc2300* | Permease AmpG | G > A | Premature stop codon |
| **Sup$_9$** | Chr1_2723906 | *vc2542* | Murein peptide ligase Mpl | TAA > TA | Premature stop codon |
| **Sup$_{10}$** | Chr1_2595642 | *vc2421* | Amidase AmpD | CGGGGGGC > CGGGGGC | Premature stop codon |
| **Sup$_{11}$** | Chr1_749620 | *vc0700* | Soluble lytic transglycosylase | TAAAAAAG > TAAAAAG | Premature stop codon |
| **Sup$_{12}$** | Chr1_2454522 | *vc2300* | Permease AmpG | GCCTTCCGGATAGATAC > GC | In frame deletion (Δ143-147) |
| **Sup$_{13}$** | Chr1_2724351 | *vc2542* | Murein peptide ligase Mpl | AGCACGTATTCCACG > AG | Premature stop codon |

[a] Accordingly to the NCBI reference sequences of *V. cholerae* N16961 strain.
[b] Nucleotide and amino acid changes are indicated for the coding or protein sequence respectively.

**Supplementary Fig. 3. Isolation of spontaneous *ΔldcV* suppressor mutants and sequencing results. a.** Emergence of spontaneous suppressor mutants in aged *ΔldcV* colonies on $LB_{10}$-agar plates. **b-c.** Phenotypic analysis of the 13 sequenced Δ*ldcV* spontaneous suppressor mutants: PG density of $LB_{10}$ stationary phase cultures (normalised to the WT) (**b**), and growth kinetics in $LB_0$ (**c**). **d-e**. Schematic representation (**d**) and list (**e**) of mutations found in the genome of sequenced spontaneous *ΔldcV* suppressors. Data represent mean ± s.d. of 2 biological replicates; **P* < 0.05, unpaired t-test.

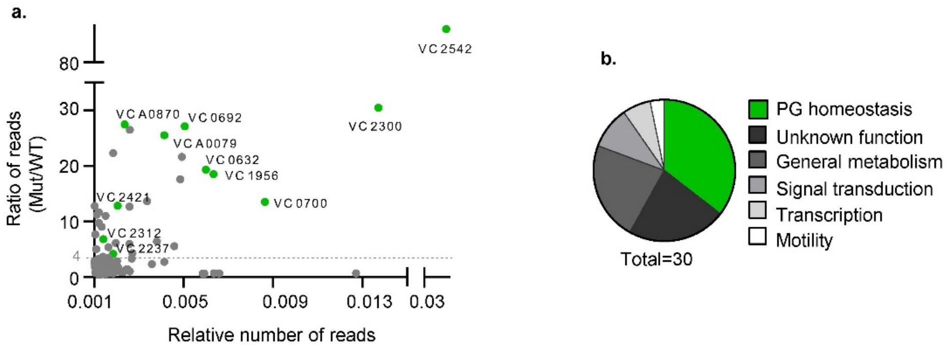


c.

| Locus | Encoded product and annotated function | Ratio reads* Mut/WT |
|---|---|---|
| ***vc2542*** | Mpl_murein peptide ligase | 87.22 |
| ***vc2300*** | AmpG_muropeptide transporter | 30.48 |
| ***vca0870*** | PbpG_D-alanyl-D-alanine endopeptidase | 27.50 |
| ***vc0692*** | NagZ_beta-hexosaminidase | 27.15 |
| ***vc1880*** | Conserved hypothetical protein transmembrane | 26.55 |
| ***vca0079*** | ShyA endopeptidase | 25.53 |
| ***vc2291*** | NqrE_NADH:ubiquinone oxidoreductase | 22.35 |
| ***vc2132*** | FliG_Flagellar motor switch protein | 21.64 |
| ***vc0632*** | DacB_D-alanyl-D-alanine carboxy/endopeptidase | 19.31 |
| ***vc1956*** | MltB_membrane-bound lytic murein transglycosylase B | 18.54 |
| ***vc2294*** | NqrB_NADH:ubiquinone oxidoreductase | 17.59 |
| ***vc2299*** | Peptidyl-prolyl cis-trans isomerase A | 13.67 |
| ***vc0700*** | Slt_Soluble lytic murein transglycosylase | 13.53 |
| ***vc2421*** | AmpD_1,6-anhydro-N-acetylmuramyl-L-alanine amidase | 12.85 |
| ***vc0441*** | Bis(5`-nucleosyl)-tetraphosphatase | 12.81 |
| ***vc1170*** | Tryptophan synthase beta subunit | 12.67 |
| **IG_*vca0869*** | Hypothetical protein | 11.65 |
| ***vc1075*** | Conserved hypothetical protein | 11.24 |
| ***vc2290*** | NADH:ubiquinone oxidoreductase, beta subunit | 10.98 |
| ***vc0758*** | Conserved hypothetical protein | 9.75 |
| **IG_*vc2472*** | Conserved hypothetical protein | 9.06 |
| ***vc1776*** | N-acetylneuraminate lyase, putative | 7.64 |
| ***vc2312*** | MltA_membrane-bound lytic murein transglycosylase A | 6.82 |
| ***vc0175*** | Deoxycytidylate deaminase-related protein | 6.42 |
| ***vc0088*** | Conserved hypothetical protein | 6.12 |
| ***vc1990*** | ATP-dependent helicase, DinG family | 5.96 |
| ***vc2215*** | Cation transport ATPase, E1-E2 family | 5.52 |
| ***vc2453*** | Sensor histidine kinase/response regulator | 5.36 |
| **IG_*vc1797*** | Hypothetical protein | 5.01 |
| ***vc2237*** | MltD_membrane-bound lytic murein transglycosylase D | 4.14 |

* normalised to the total number of reads

**Supplementary Fig. 4. Tn-seq of *ΔldcV* suppressor mutants selected on LB$_{0}$. a.** Genomic distribution of the transposon-insertions obtained from sequencing the *ΔldcV*-LB$_0$ Tn-library, represented based on the relative number of reads per gene (X axis) and on the fold change of these values when compared with the reads of an WT-LB$_0$ library (Y axis). In green are highlighted genes annotated to be involved in PG homeostasis. **b-c.** Functional classification of those loci found to be at least 4 times more represented (mutated) in the *ΔldcV*-LB$_0$ Tn-library than in the control WT-LB$_0$ (**b**) and their identity (**c**).

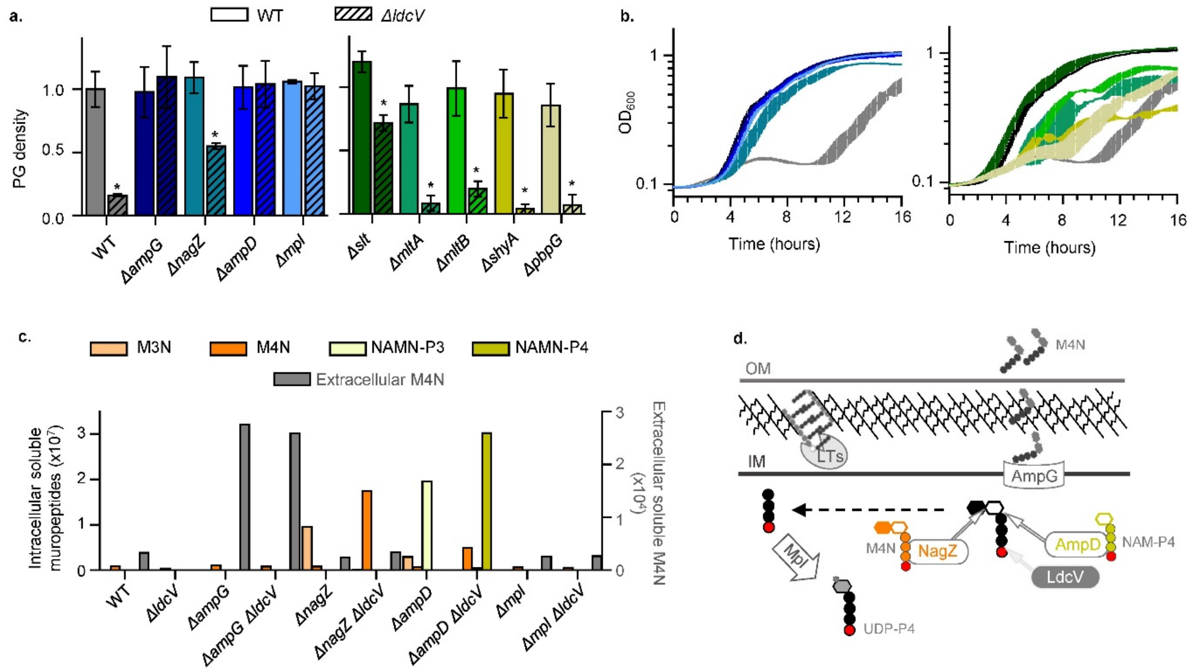


**Supplementary Fig. 5. Phenotypic analysis *ΔldcV* suppressor mutants. a-b.** Quantification of the amount of PG per cell and (normalized to the WT) (**a**) and growth kinetics in $LB_0$ of the WT, *ΔldcV,* and different PG-recycling single and *ΔldcV*-double mutants (**b**). **c-d.** Level of soluble muropeptides detected by $MS^{e}$ in the cytosol (different colours, left axis) and in the extracellular medium (grey, right axis) of indicated mutant strains (**c**); and schematic representation for explaining the accumulation of these forms in the different PG-recycling mutants indicating in red the D-Ala at fourth position maintained in the *ΔldcV* background leading to accumulation of tetrapeptide forms (**d**). Anhydro-murotripeptide (M3N), anhydro-murotetrapeptide (M4N), anhydro-muramyl tripeptide (NAMN-P3) anhydro-muramyl tetrapeptide (NAMN-P4). Data in a represent mean ± s.d. of at least 2 biological replicates; *$P < 0.05$, unpaired t-test.

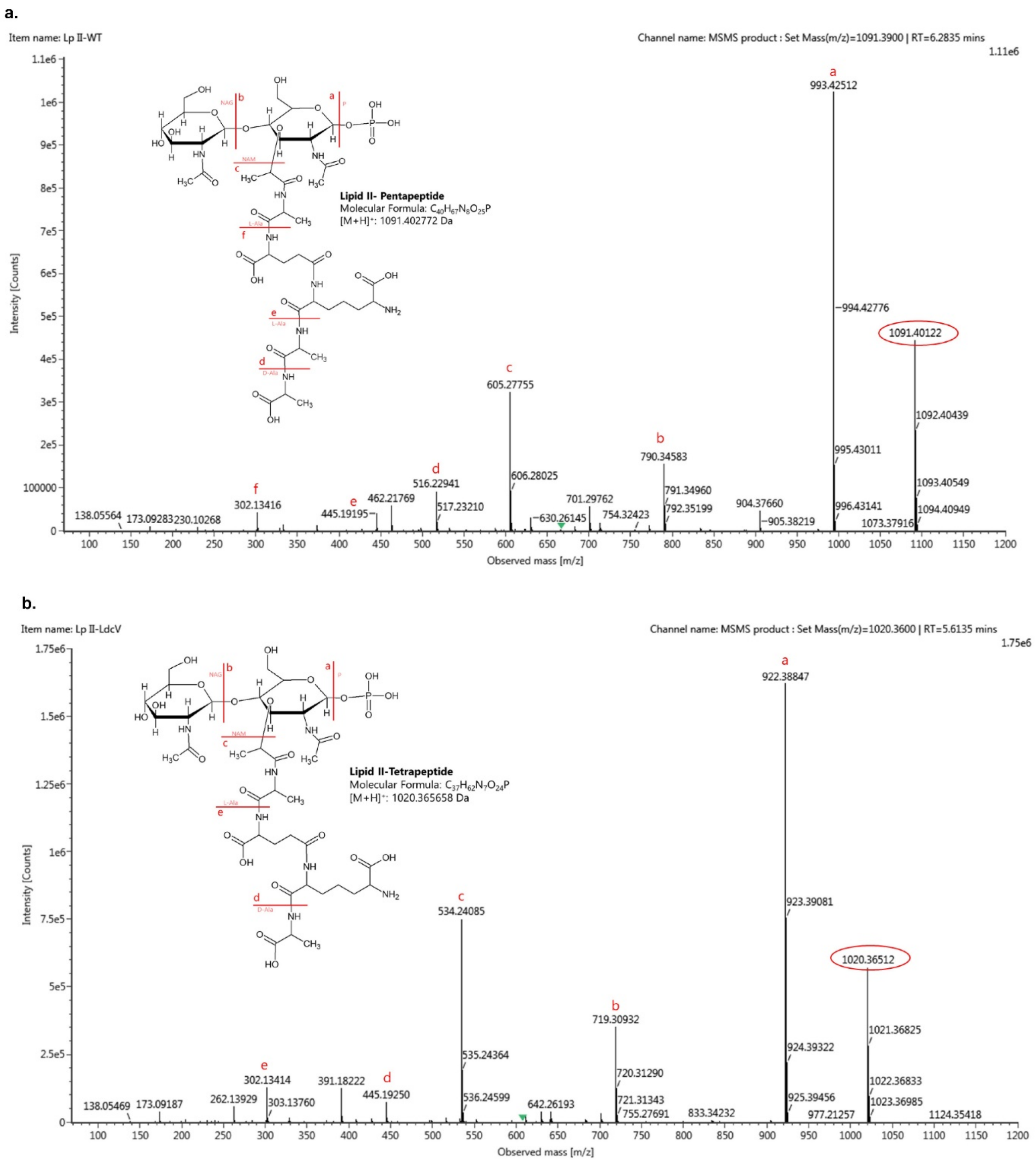


**Supplementary Fig. 6. Detection of delipidated Lipid II forms.** MS/MS profile of delipidated Lipid II-P5 (**a**) and Lipid II-P4 (**b**) detected by targeted MS in the Lipid-II sample-preparations obtained from WT and *ΔldcV* cultures respectively.

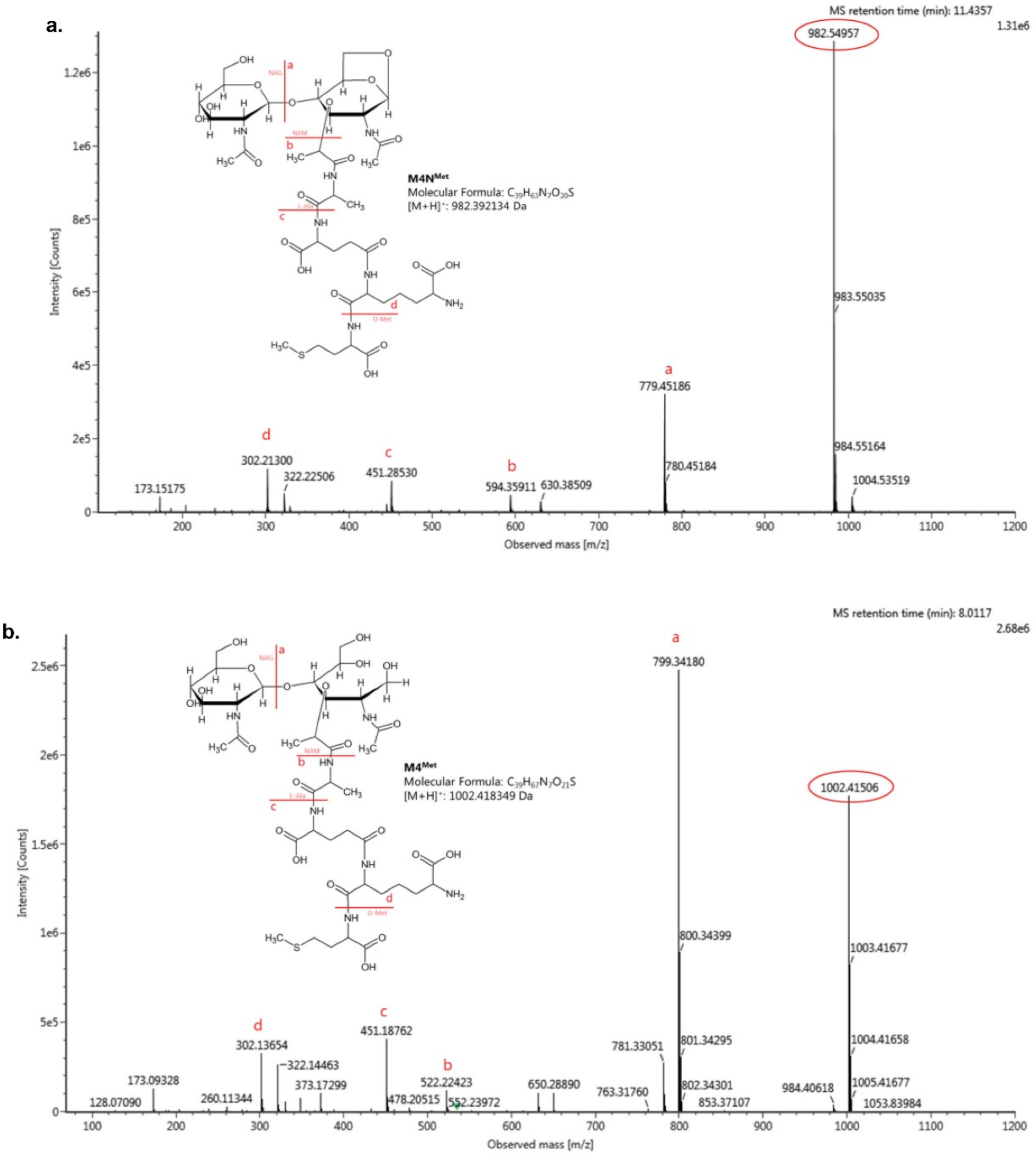


**Supplementary Fig. 7.** Assay to detect incorporation of recycled D-Met-modified tetrapeptides into the PG. MS/MS profile of: purified M4N^Met used for treatment of *ΔldcV* cultures (**a**); and M4^Met produced by *in vitro* reactions with LdtA to use as standard (**b**).

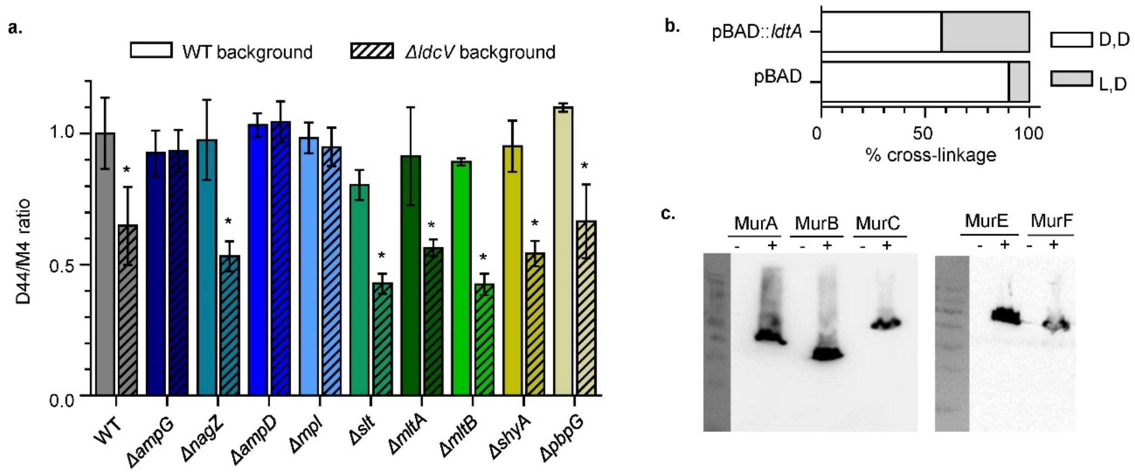


**Supplementary Fig. 8. a.** D44/M4 ratio (normalised to the value obtained for the WT) calculated for the PG of $LB_{10}$ stationary cultures of different single and double mutants. **b.** Percentage of D,D or L,D cross-linkage (respective to the total amount of PG cross-linkage) of *ΔldcV* mutant carrying the $pBAD_{18}$ empty plasmid or the pBAD::*ldtA* construction growing in $LB_{10}$ with 0.2% of arabinose. **c.** Western blot showing the amount of several Mur-proteins in arabinose-induced cultures (+) of *V. cholerae* carrying specified $pBAD_{18}$::*mur*-6xHis construction. Data represent mean ± s.d. of at least 3 biological replicates; **P* < 0.05, unpaired t-test.

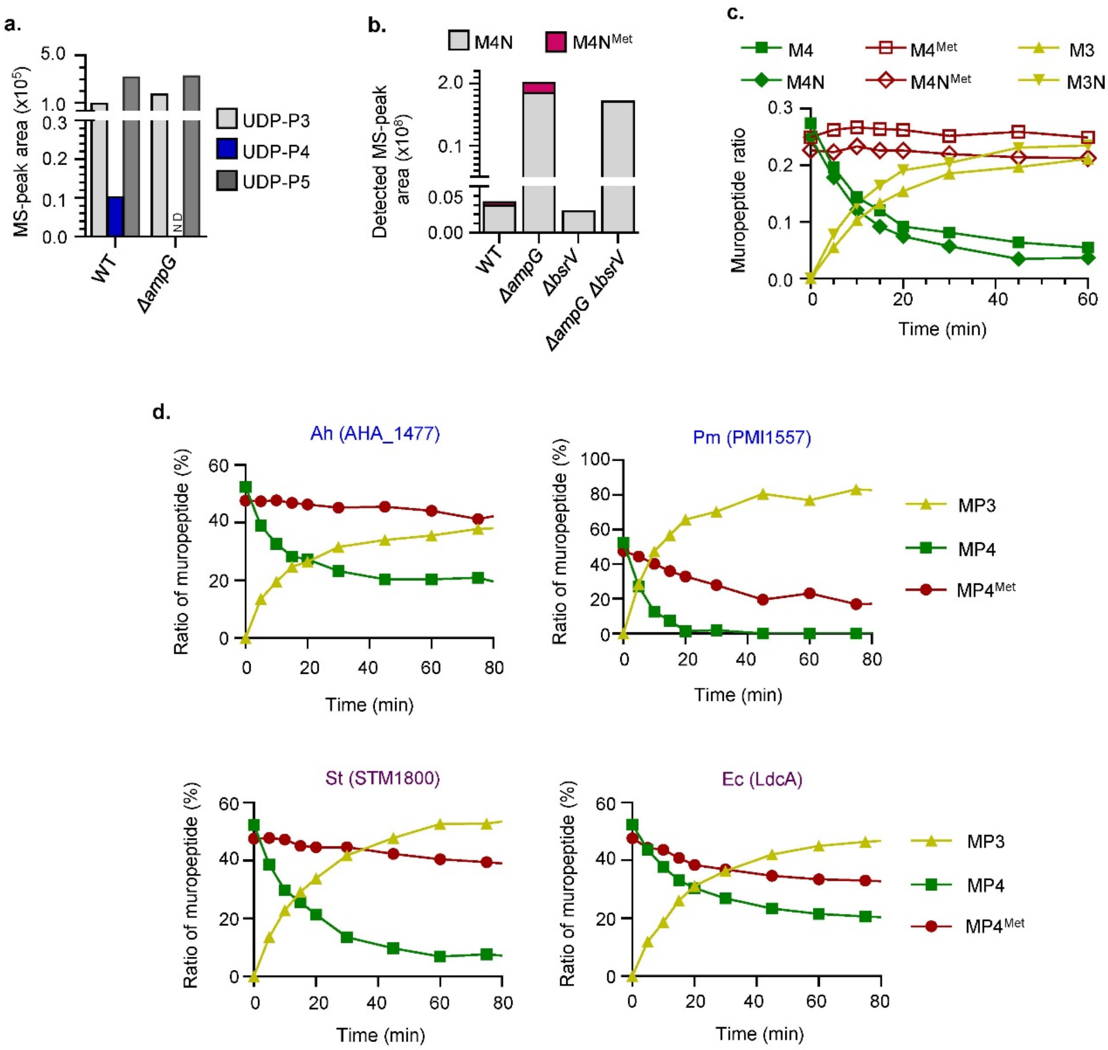


**Supplementary Fig. 9. a.** Cytosolic UDP-forms detected by targeted MS in exponential phase cultures of indicated strains growing in LB+L-Met. **b.** Extracellular M4N$^{Met}$ detected in stationary cultures of indicated strains growing in LB supplemented with L-Met. **c.** Dynamics of substrate consumption of LdcV (M4: murotetrapeptide; M4$^{Met}$: murotetrapeptide with D-Met at fourth position; M4N: anhydro-murotetrapeptide; M4N$^{Met}$: anhydro-murotetrapeptide with D-Met at fourth position) and product production (M3: murotripeptide; M3N: anhydro-murotripeptide). **d**. Dynamics of substrate consumption during *in vitro* reactions of purified L,D-carboxypeptidases from *Aeromonas hydrophila* (Ah), *Proteus mirabilis* (Pm) *Salmonella enterica* serovar Typhimurium (St) and *E. coli* (Ec) (MP4= M4 + M4N; MP4$^{Met}$= M4$^{Met}$ + M4N$^{Met}$ ; MP3= M3 + M3N). ND: not detected.

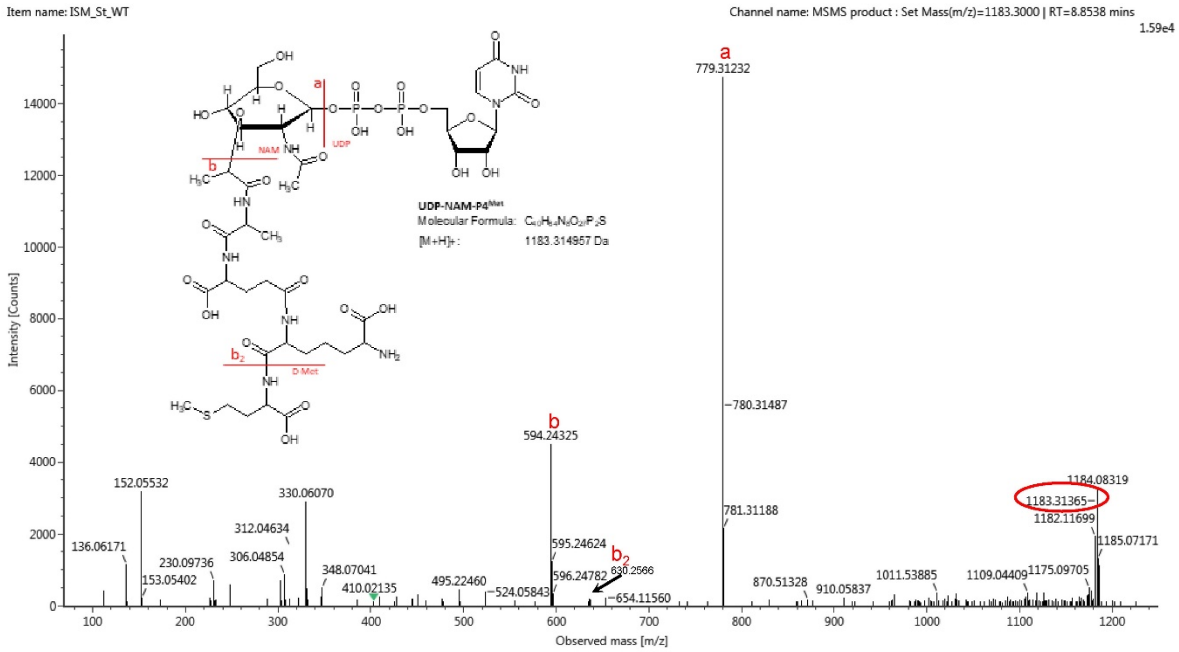


**Supplementary Fig. 10. Detection of D-Met modified tetrapeptide precursors. a.** MS/MS profile of UDP-P4$^{Met}$ detected by targeted MS in the cytosolic pool of PG-precursors of the WT strain growing until stationary phase in LB + L-Met.

**Supplementary Table 1. Bacterial strains.**

| ***Vibrio cholerae* N16961 derivative strains:** | **FC_Number** | **Source/Ref** |
|---|---|---|
| Δ*mrcA* | FC2388 | Lam *et al.* 2009 |
| Δ*ldcV* (*vc2153*) | FC2029 | this study |
| Δ*ampD* (*vc2421*) | FC2505 | this study |
| Δ*ampG* (*vc2300*) | FC2506 | this study |
| Δ*nagZ* (*vc0692*) | FC2507 | this study |
| Δ*mpl* (*vc2542*) | FC2508 | this study |
| Δ*slt* (*vc0700*) | FC2509 | this study |
| Δ*mltA* (*vc2321*) | FC2652 | this study |
| Δ*mltB* (*vc1956*) | FC2656 | this study |
| Δ*shyA* (*vca0079*) | FC2654 | this study |
| Δ*pbpG* (*vca0870*) | FC2658 | this study |
| Δ*ampD* Δ*ldcV* | FC2510 | this study |
| Δ*ampG* Δ*ldcV* | FC2511 | this study |
| Δ*nagZ* Δ*ldcV* | FC2512 | this study |
| Δ*mpl* Δ*ldcV* | FC2513 | this study |
| Δ*slt* Δ*ldcV* | FC2514 | this study |
| Δ*mltA* Δ*ldcV* | FC2653 | this study |
| Δ*mltB* Δ*ldcV* | FC2657 | this study |
| Δ*shyA* Δ*ldcV* | FC2655 | this study |
| Δ*pbpG* Δ*ldcV* | FC2659 | this study |
| *Δslt ΔampG* | FC2766 | this study |
| *ΔmltA ΔampG* | FC2767 | this study |
| *ΔmltB ΔampG* | FC2768 | this study |
| *ΔshyA ΔampG* | FC2769 | this study |
| *ΔpbpG ΔampG* | FC2770 | this study |
| *ΔldtA ΔldtB ΔbsrV* | FC2770 | Cava *et al*. 2011 |
| *ΔldtA ΔldtB ΔbsrV ΔampG* | FC2678 | this study |
| *ΔldtA ΔldtB ΔbsrV ΔldcV* | FC2679 | this study |
| *ΔldtA ΔldtB ΔbsrV ΔampG ΔldcV* | FC2680 | this study |
| *ΔldtA* | FC45 | Cava *et al.* 2011 |
| *ΔldtA* Δ*ldcV* | FC2765 | this study |
| ***Escherichia coli* strains** | **FC_Number** | **Source/Ref** |
| BL21 pET28b::*vc2153* | FC658 | this study |
| BL21 pET28b::*ldcA* | FC2026 | this study |
| BL21 pET22b::*vca0337* | FC2665 | this study |
| BL21 pET28b::*slt70* (*E. coli*) | FC1860 | Espaillat *et al.* 2016 |
| BL21 pET28b::*ldtA* | FC1168 | Cava *et al.* 2011 |
| BL21 pET28b::AHA1477-His | FC2775 | this study |
| BL21 pET28b::PMI1557-His | FC2776 | this study |
| BL21 pET28b::STM1800-His | FC2777 | this study |

**Supplementary Table 2: Primers used for constructing deletion mutants.**

| Name | Sequence (5’→3’) | FCP number |
|---|---|---|
| vc2153-P1-XbaI-FOR | AAATCTAGAtgattagccgccaatccgttac | FCP941 |
| vc2153-P2-dapE_tailNotI-REV | TTTTTTGCGGCCGCTTTTTTcggactcacttgcggatgcact | FCP1041 |
| vc2153-P3-tailNotI-FOR | AAAAAAGCGGCCGCAAAAAAggtataacatggagatactgtt | FCP943 |
| vc2153-P4-SalI-REV | AAAGTCGACatgttgcgccaagatgaagcg | FCP944 |
| vc2153 E1 | GGGCGCACAAGTGGTTGAGCTCGGTCCCG | FCP303 |
| vc2153 E2 | GCCCAAATTTCATGTTGGCGGTG | FCP302 |
| vc2421-P1-XbaI-FOR | AAATCTAGAcgctgtgctaatcgcctgacg | FCP1060 |
| vc2421-P2-tailNotI-REV | TTTTTTGCGGCCGCTTTTTTccttactccttgcttatatct | FCP1061 |
| vc2421-P3-tailNotI-FOR | AAAAAAGCGGCCGCAAAAAAataggttttagttggtgcgaa | FCP1062 |
| vc2421-P4-XbaI-REV | AAATCTAGAgcgctcaaaccacatggtatc | FCP1063 |
| vc2421 E1 | tcatcgcaagtcgaatgtcc | FCP1059 |
| vc2421 E2 | ctgaccgagtatcacgatgc | FCP1064 |
| vc2300-P1-XbaI-FOR | AAATCTAGAcgcaatatgtggctttagtcg | FCP1072 |
| vc2300-P2-tailNotI-REV | TTTTTTGCGGCCGCTTTTTTaacgagtccttttgcatgtgat | FCP1073 |
| vc2300-P3-tailNotI-FOR | AAAAAAGCGGCCGCAAAAAAgtaaggcgcacaccattcacc | FCP1074 |
| vc2300-P4-XbaI-REV | AAATCTAGAgagcaggtgatgaaatttctc | FCP1075 |
| vc2300 E1 | aggtcgaatgcagaacgttc | FCP1071 |
| vc2300 E2 | ccgtattgactgagcctatg | FCP1076 |
| vc0692-P1-XbaI-FOR | AAATCTAGAgtgcgacttgctcaacaatgg | FCP1066 |
| vc0692-P2-tailNotI-REV | TTTTTTGCGGCCGCTTTTTTgaccaaacagaatcacaccac | FCP1067 |
| vc0692-P3-tailNotI-FOR | AAAAAAGCGGCCGCAAAAAAatagtttcctcatagaagta | FCP1068 |
| vc0692-P4-XbaI-REV | AAATCTAGAgtgaagattatgatggtttag | FCP1069 |
| vc0692 E1 | cacaccatccaaactggttc | FCP1065 |
| vc0692 E2 | gtgaagattatgatggtttag | FCP1070 |
| vc2542-P1-XbaI-FOR | AAATCTAGAcacatcgatattggaagagc | FCP2077 |
| vc2542-P2-tailNotI-REV | TTTTTTGCGGCCGCTTTTTTgccgcccatgaacgtgcc | FCP2078 |
| vc2542-P3-tailNotI-FOR | AAAAAAGCGGCCGCAAAAAAgctcgcggcactgcaacag | FCP2079 |
| vc2542-P4-XbaI-REV | AAATCTAGAttgatcatagccccaacgcg | FCP2080 |
| vc2542 E1 | Gacctaaggttagtgaagtg | FCP2075 |
| vc2542 E2 | gtaggtacacctgatagtccg | FCP2076 |
| vc0700-P1-XbaI-FOR | AAATCTAGAttgtcaatgcctgccatg | FCP1777 |
| vc0700-P2-tailNotI-REV | TTTTTTGCGGCCGCTTTTTTcatgagatctcatcc | FCP1778 |
| vc0700-P3-tailNotI-FOR | AAAAAAGCGGCCGCAAAAAAcgtacaaggccgcttc | FCP1779 |
| vc0700-P4-XbaI-REV | AAATCTAGAcatctcacccgcagc | FCP1780 |
| vc0700 E1 | cgaactctttgcactcagcc | FCP1776 |
| vc0700 E2 | tcgagcgactaatgcttcac | FCP1781 |
| vca0079 E1 | Acaactaagttgctgccagc | FCP1770 |
| vca0079-P1-XbaI-FOR | AAATCTAGAgatggagtttgacatg | FCP1771 |
| vca0079-P2-tailNotI-REV | TTTTTTGCGGCCGCTTTTTTgatatcaccagtttacctg | FCP1772 |
| vca0079-P3-tailNotI-FOR | AAAAAAGCGGCCGCAAAAAAcgcaataaatcgagtatg | FCP1773 |
| vca0079-P4-XbaI-REV | AAATCTAGAcgatgagctgacgcattc | FCP1774 |
| vca0079 E2 | tctcaacgcttgcctcaatc | FCP1775 |
| vc2312-XbaI-P1fw | AAAATCTAGACAAGAGGCTGGATAAAGT | FCP1907 |
| vc2312-P2rev | ATCAGGAAAAAAAAAGAGGTCCATGGTT | FCP1908 |
| vc2312-P3fw | GGACCTCTTTTTTTTTCCTGATATTGTGC | FCP1909 |
| vc2312-XbaI-P4rev | AAAATCTAGAGTAAGCCAGAAGAGAAGC | FCP1910 |
| vca0870-P1-SalI-FOR | AAAGTCGACgtgattgggatatcaccc | FCP2647 |
| vca0870-P2-tailNotI-REV | TTTTTTGCGGCCGCTTTTTTttttgaattcactgaacg | FCP2648 |
| vca0870-P3-tailNotI-FOR | AAAAAAGCGGCCGCAAAAAAtgacgatccaatctcactc | FCP2649 |
| vca0870-P4-SalI-REV | AAAGTCGACtggaatgatctcggggtc | FCP2650 |
| vca0870-E1 | ccagaccgtataccaaaatc | FCP2646 |
| vca0870-E2 | actagggtgagattggtgag | FCP2651 |
| vc1956-P1-XbaI-FOR | AAATCTAGActctatgccttacggagag | FCP2653 |
| vc1956-P2-tailNotI-REV | TTTTTTGCGGCCGCTTTTTTagatgccctccttgagcc | FCP2654 |
| vc1956-P3-tailNotI-FOR | AAAAAAGCGGCCGCAAAAAAtaacctgttggattggcg | FCP2655 |
| vc1956-P4-XbaI-REV | AAATCTAGActtcgacgttcaactcag | FCP2656 |
| vc1956-E1 | tgctggtcaatcttgagcag | FCP2652 |
| vc1956-E2 | aacagagggctcaatgagag | FCP2657 |

**Supplementary Table 3: Primers used to construct the plasmids for complementation and overexpression experiments.**

| Name | Sequence (5'→3') | FCP number |
|---|---|---|
| ldtA-EcoRI-RBS-FOR | atgcGAATTCaggaggggcgttttgatgaaaagc | FCP2499 |
| ldtAhis-XbaI-REV | gcatTCTAGATTAgtgatgatgatgatgatgaattttagtcaaatttgg | FCP2563 |
| murA-EcoRI-RBS-FOR | atgcGAATTCaggaggcctttaaggtttttatgg | FCP2962 |
| murAhis-XbaI-REV | gcatTCTAGATTAgtggtggtggtggtggtggtcgcggaaacgctcaat | FCP2963 |
| murB-EcoRI-RBS-FOR | atgcGAATTCaggagggaggtagaatatggccag | FCP2964 |
| murBhis-XbaI-REV | gcatTCTAGATTAgtggtggtggtggtggtgtgcttgttcactcatcca | FCP2965 |
| murC-EcoRI-RBS-FOR | atgcGAATTCaggaggcgagaacagaattatgac | FCP2966 |
| murChis-XbaI-REV | gcatTCTAGATTAgtggtggtggtggtggtgaatctgttgcatacgccc | FCP2967 |
| murE-EcoRI-RBS-FOR | atgcGAATTCaggaggcaaatcctcaacattgctc | FCP3137 |
| murEhis-XbaI-REV | gcatTCTAGATTAgtggtggtggtggtggtgtggttgtaactccagtaa | FCP3138 |
| murF-EcoRI-RBS-FOR | atgcGAATTCaggagggagttacaaccatgattc | FCP3139 |
| murFhis-XbaI-REV | gcatTCTAGATTAgtggtggtggtggtggtgtgcaaatttctccttgag | FCP3140 |

**Supplementary Table 4: Primers used to construct the plasmids for protein overexpression and purification.**

| Name | Sequence (5'→3') | FCP number |
|---|---|---|
| vc2153_Nco_FOR | AAACCATGGGGgtgtgtacgtattgcggatttagagaaattgac | FCP269 |
| vc2153_his-Eco-REV | AAAGAATTCttagtggtggtggtggtggtgtacctcacaaatattcgtgatgaagcgc | FCP268 |
| ldcA-NcoI-FOR | atgcccatgggctctctgtttcacttaattgccccatcgggtta | FCP937 |
| ldcA_His-EcoRI-REV | atgcgaattcttaGtggtggtggtggtggtgcattttaagaacaggatgac | FCP938 |
| vca0337-NdeI-FOR | TGCCATATGttatatgccaaagctttaag | FCP1418 |
| vca0337_His-NotI-REV | AAAGCGGCCGCttttgcttttacccaacgatc | FCP1419 |
| AHA1477-NcoI-FOR | atgcccatgggcacccaggagcaactgctggggt | FCP2942 |
| AHA1477-EcoRI-His-REV | atgcgaattcttagtggtggtggtggtggtgtgtgctactcctttcgtcgg | FCP2943 |
| PMI1557-NcoI-FOR | atgcccatgggcataaccccctttaatgttaacgg | FCP2944 |
| PMI1557-EcoRI-His-REV | atgcgaattcttagtggtggtggtggtggtgttctggtagcacaatatagc | FCP2945 |
| STM1800-NcoI-FOR | atgcccatgggcgaattaccatgtctctgtttca | FCP2946 |
| STM1800-EcoRI-His-REV | atgcgaattcttagtggtggtggtggtggtgcaattgcagcgtaggatggc | FCP2947 |
